# The intersectional genetics landscape for human

**DOI:** 10.1101/552984

**Authors:** Andre Macedo, Alisson M. Gontijo

## Abstract

The human body is made up of hundreds, perhaps thousands of cell types and states, most of which are currently inaccessible genetically. Genetic accessibility carries significant diagnostic and therapeutic potential by allowing the selective delivery of genetic messages or cures to cells. Research in model organisms has shown that single regulatory element (RE) activities are seldom cell type specific, limiting their usage in genetic systems designed to restrict gene expression posteriorly to their delivery to cells. Intersectional genetic approaches can theoretically increase the number of genetically accessible cells, but the scope and safety of these approaches to human have not been systematically assessed due primarily to the lack of suitable thorough RE activity databases and methods to explore them. A typical intersectional method acts like an AND logic gate by converting the input of two or more active REs into a single synthetic output, which becomes unique for that cell. Here, we systematically assessed the intersectional genetics landscape of human using a curated subset of cells from a large RE usage atlas obtained by Cap Analysis of Gene Expression sequencing (CAGE-seq) of thousands of primary and cancer cells (the FANTOM5 consortium atlas). We developed the heuristics and algorithms to retrieve AND gate intersections and quality-rank them intra- and interindividually. We find that >90% of the 154 primary cell types surveyed can be distinguished from each other with as little as 3 to 4 active REs, with quantifiable safety and robustness. We call these minimal intersections of active REs with cell-type diagnostic potential “Versatile Entry Codes” (VEnCodes). Each of the 158 cancer cell types surveyed could also be distinguished from the healthy primary cell types with small VEnCodes, most of which were highly robust to intra- and interindividual variation. Finally, we provide methods for the cross-validation of CAGE-seq-derived VEnCodes and for the extraction of VEnCodes from pooled single cell sequencing data. Our work provides a systematic view of the intersectional genetics landscape in human and demonstrates the potential of these approaches for future gene delivery technologies in human.

## INTRODUCTION

The exact number of different cell types that make up the body of a human adult is yet to be defined, but is expected to be in the order of several hundred, perhaps thousands of different cell types (Valentine *et al.*, 1994; Carrol, 2001). Major efforts have recently been launched to attempt to catalogue and molecularly describe every cell type in different tissues of the human body (Andersson *et al.*, 2014; FANTOM Consortium and the RIKEN PMI and CLST (DGT), 2014; Macosko *et al.*, 2015; Bahar Halpern *et al.*, 2017; Regev *et al.*, 2017; Hon *et al.*, 2018). The number and the complexity of cell types increases further when one considers that cells exist in different states, not only when a cell divides or undergoes successive differentiation steps during normal developmental processes, but also when a cell becomes infected, cancerous or specifically responds to physical or chemical stimuli (Valentine *et al.*, 1994; Carrol, 2001; Macosko *et al.*, 2015).

A major challenge in biology and biomedicine has been to genetically identify and deliver genetically encoded messages to a specific cellular type and/or state within complex organisms. Most gene delivery systems are limited by the technology available to distinguish the desired cellular types and/or states between themselves prior to gene delivery; most technologies relying primarily on cell-surface markers for selectivity (Lukashev and Zamyatnin, 2016; Hardee *et al.*, 2017). These markers are seldom cell-specific, and this lack of specificity inevitably leads to DNA delivery to unwanted cells. This can have negative consequences, such as introducing undesired artefacts in research studies or side-effects in gene-therapy-based interventions. Additionally, the usage of sporadically defined cell surface markers for cellular targeting restricts both the ability to systematize the generation of cell-specific gene delivery vectors and to scale this system up for any cell type or state in any organism.

An alternative to these “pre-DNA delivery” selectivity procedures is to use cell-type- and cell-state-unspecific viral or non-viral DNA delivery systems (Duan, 2016; Wong *et al.*, 2016), and work out the cell specificity post-delivery by exploring unique genetic properties of the target cell. The transcriptional program of any given cell reflects, at the most basic level, a unique combination of binary on/off states of the regulatory elements (REs) present in the genome. REs can be used multiple times by different cells either at different anatomical sites, time points of life history or during disease or environmental responses (Mallo, 2006; Luan *et* al., 2006; ENCODE Project Consortium, 2012; Mortazavi *et al.*, 2013; Kron *et al.*, 2014; Andersson *et al.*, 2014; FANTOM Consortium and the RIKEN PMI and CLST (DGT), 2014). Therefore, while the activity of a single carefully-chosen RE could theoretically provide sufficient specificity to identify a particular cell type and/or state post-DNA delivery in some cases, it is unlikely to provide the required specificity to distinguish most cell types and/or states between themselves (Mallo, 2006; Luan *et al.*, 2006).

Aware of this fact, developmental biologists studying model organisms have devised intersectional genetic methods to increase target cell specificity of gene drivers by exploring the anatomical overlap between expression patterns driven by two independent REs (Lakso *et al.*, 1992; Struhl and Basler, 1994; Awatramani *et al.*, 2003; Suster *et al.*, 2004; Stockinger *et al.*, 2005; Luan *et al.*, 2006; Farago *et al.*, 2006). Similarly, molecular and synthetic biologists have engineered systems that use Boolean logic to sense different cell states in bacteria and yeast (Siuti *et al.*, 2013, Nissim *et al.*, 2007). In many of these synthetic computational systems, the REs are the inputs which will pass through a typical AND gate and give a single genetically-defined output (Figure 1). Similar systems have been applied to mammalian cells, where they are able to distinguish between different cancer cell types or detect cancer cells arising from normal cells in vitro (Nissim and Bar Ziv, 2010; Liu *et al.*, 2014; Morel *et al.*, 2016). Despite being successful, the full potential of this type of intersectional approach has never been evaluated or applied systematically to generate drivers for every cell type in a body, even less so to a complex organism like human, which lacks thoroughly developmentally-characterized gene drivers.

**Figure 1.**
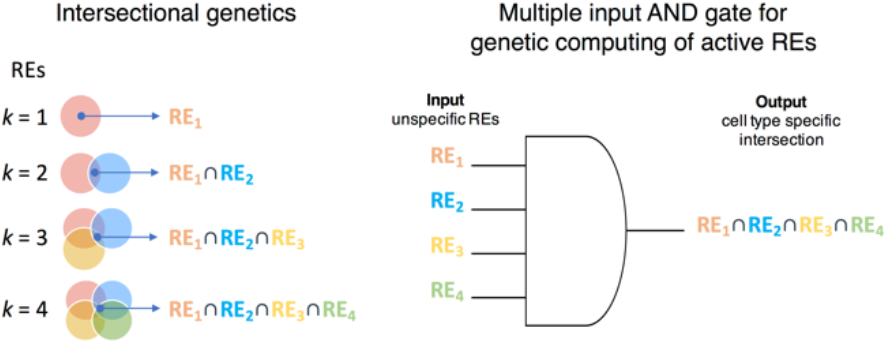
Intersectional genetics. Scheme of the intersectional genetics approach to obtain cell-type specific drivers by restricting expression to the cells where two or more REs with broader activity overlap (intersect). REs are the inputs that will pass through a typical AND logic gate and give a single genetically-defined output in the cells where the RE activities intersect.

Here, we hypothesized that the majority of cell types and/or cell states in human could be distinguished post-DNA delivery using multiple input AND gates (intersectional methods of active REs, Figure 1), and that the intersecting inputs could be obtained, quality-ranked, and cross-validated using currently publicly available RE usage databases.

## RESULTS

### Data preparation / normalization

To quantify how cellular specificity scales with the number of intersecting active REs (*k*), we developed algorithms and scripts using Python language to analyze genome-wide data on promoter and enhancer usage for hundreds of primary human cell types obtained by the FANTOM5 consortium (Andersson *et al.*, 2014; FANTOM Consortium and the RIKEN PMI and CLST (DGT), 2014; Lizio *et al.*, 2015). Briefly, the FANTOM5 data consists of curated subsets of transcriptional start site “peaks” determined by capped analyses of gene expression (CAGE)-sequencing (CAGE-seq). The height of each CAGE-seq peak provides quantitative information in normalized tags per million (TPM) values, which is interpreted as being directly proportional to the activity of the promoter or enhancer that it represents.

Before analyzing the FANTOM5 data, we manually curated the FANTOM5 human cell type database consisting of 184 distinct cell types from multiple donors (giving a total of 562 datasets), by selecting for healthy primary cells and removing cell treatments/infections and cells obtained from cancer samples (Figure S1). We also attempted to remove datasets that were less likely to represent single cell types. Examples of the samples removed during curation are: datasets from cells infected with Salmonella or Candida albicans, datasets for cells labeled “whole blood”, and datasets from mesenchymal precursor cells obtained from cancer samples. Some datasets were merged into a single cell type category, for example: “CD8+ T Cells (pluriselect)” and “CD8+ T Cells”, and “Melanocyte dark” and “Melanocyte light” were treated as single cell type categories, respectively. This curation resulted in a list of 154 distinct primary cell types from multiple donors, giving a total of 537 samples and averaging ~3.5 samples (donors) per cell type (range 2-6). Table S1 contains the list of curated cell types used in this study as well as all of the excluded and merged categories.

The total number of possible RE combinations for a target cell type is *C(*r, k*) = (r!/(k!(r - k)!)*, where *r* stands for the number of REs of the database (*e.g*., 201 802 promoters in FANTOM5), and *k* for the number of REs chosen to combine. For *k* = 4, this gives 6.9 x 10^19^ possible combinations. To ask if any combination is specific for the target cell type, however, we need to ask if the *k* combined elements are all active in the given cell type AND at least one of the *k* elements is inactive in each of the other cell types in the database. If the *k* elements could be binarized into active (TRUE) an inactive (FALSE) categories, this question can be asked using boolean logic gate functions such as: *(^1^k_1_ AND ^1^k_2_ AND … ^1^k_n_) AND ((^2^k_1_ AND ^2^k_2_ AND…. ^2^k_n_) NOR (^3^k_1_ AND ^3^k_2_… AND ^3^k_n_)… NOR (^n^k_1_ AND ^n^k_2_… AND ^n^k_n_))*, where 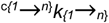 represents the status of the RE element *k* in cell type *c* (where the target cell type is 1). The truth table for this function has 2^(c*k)^ rows, which for 154 cell types and *k* = 4 gives 2.7 x 10^185^ rows. Clearly, saturating the search for all possible combinations for any given cell type and testing them by brute-force is a daunting computational task.

The complexity of the database for a given cell type can nevertheless be reduced for each search using heuristic methods. For instance, REs that are inactive in the target cell or active in the target cell and also active in most other non-target tissues (*e.g.*, REs of housekeeping genes) are not helpful for the purpose of making cell-type-specific intersectional gene drivers.

Hence, to increase the likelihood of finding fruitful intersections and to reduce database complexity and computing time, we applied several filters on the database to select for sparsely-active REs. The first step is to define RE activity thresholds. We decided to be conservative and apply different activity thresholds for the target cell type and for the non-target cell types. This would increase the chances that the selected REs are truly active in the target cell type and inactive in the non-target cell type. To reduce database size and concentrate on potentially active REs in the target cell type, we created subsets of data for each target cell type where we retained only the REs that were consistently potentially ON (>0 TPM) in all donors for that cell type (Figure 2A). We next collapsed the data from multiple donors of the non-target cell types to a single non-target cell-type datapoint by averaging the expression of the multiple donors (Figure 2B). This reduces the database complexity by a factor of ~3.5.

**Figure 2.**
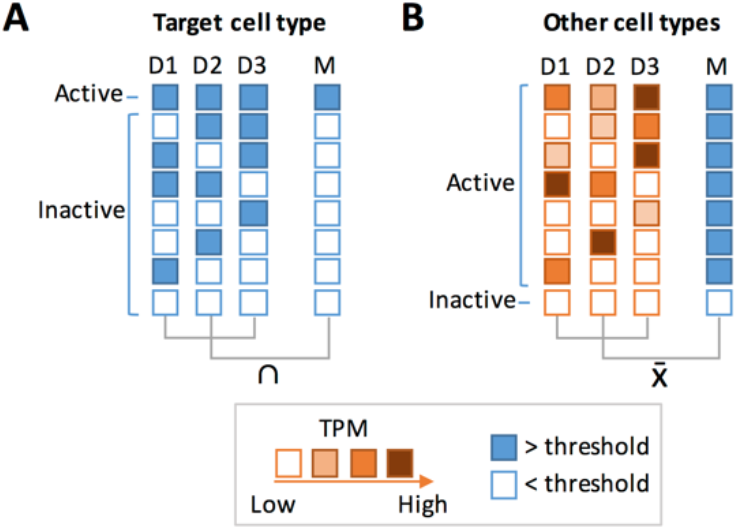
Conservative criteria for RE activity. Different conservative criteria for RE activity were applied to target **(A)** and non-target cells (“Other cell types”) **(B)**. Each row represents a possible RE activity scenario. Each box represents the activity of the RE per donor (D) or the collapsed intersection or average (M), according to the color key. REs from target cells were considered active if the intersection of all cell donors were above a TPM threshold (blue squares). REs from other cell types were considered active if the average raw TPM (M) of all donors was above the threshold.

To select for sparsely-active REs, we studied the RE activity landscape by testing the following thresholds for RE activity in the target cell type: 0.5, 1, and 2 TPM. The higher the RE activity threshold, the more stringent the RE selection is. For inactivity, we tested 0, 0.01, and 0.1 TPM in non-target cells. By applying these thresholds, we transform the continuous CAGE-seq peak data into binary datasets.

We then wrote a program that randomly samples the filtered RE landscape by choosing a combination of *k* “active” REs for a target cell type and asking whether this combination is exclusive to the target cell type compared to the other cell types of the database. We call this the “Sampling Method” (Figure 3A).

**Figure 3.**
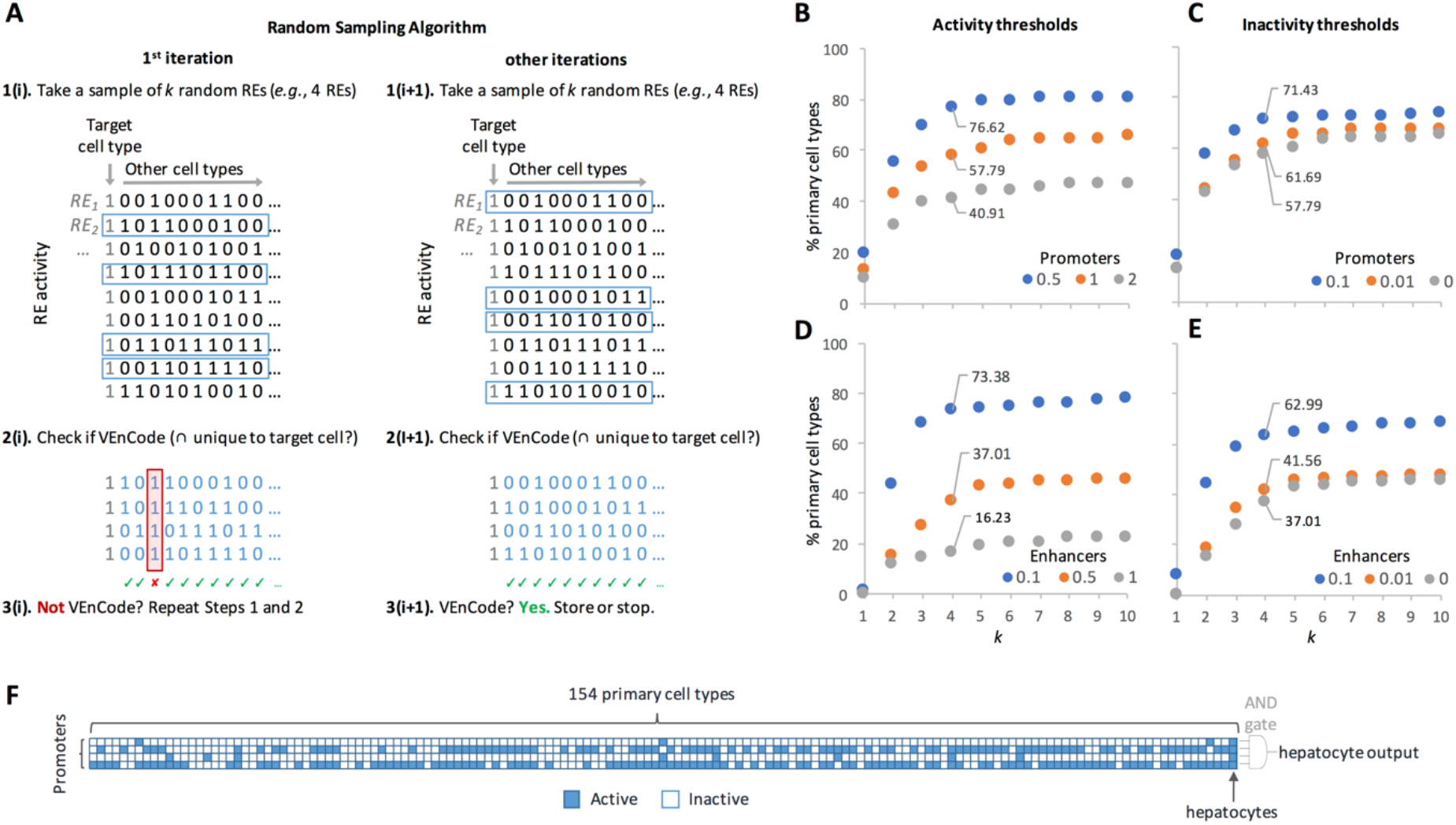
Random sampling method to find intersecting active REs (VEnCodes). **A.** Rationale for the sampling method. First, *k* REs are randomly selected from the set of REs that are active (“1”) in the target cell type. Inactive REs are depicted as “0”. Then, we ask if at least one sampled RE is inactive in each other cell type in the data set. If yes, these *k* REs satisfy VEnCode criteria or the target cell type (*e.g.*, the *k* REs must intersect exclusively in the target cell). If not, we repeat steps 1 and 2. If in the first or second iteration (i+1), the *k* REs satisfy VEnCode criteria, then the *k* RE selection is counted as a VEnCode and is stored. **B-E.** Probing the intersection genetics landscape for promoter **(B, C)** and enhancer **(D, E)** datasets using the sampling method. Plotted are the percentages of cell types found to have at least one VEnCode per *k* and different activity **(B,D)** and inactivity **(C,E)** TPM thresholds. For the activity panels, the inactivity threshold was fixed at 0 for both promoters and enhancers. For the inactivity panels, the activity thresholds were fixed at 0.5 and 0.1 for promoters and enhancers, respectively. **F.** Visual representation of a VEnCode for hepatocytes. Binary heatmap where each column represents one of the 154 primary human cell types and each row one RE from the promoter data set. Blue (active RE), white (inactive RE).

To further reduce computing time, the algorithm first selects for sparsely-active REs by removing all REs that are active in more than X% of the cell types. This removes broadly expressed REs. We start with X = 90%, but decrement 5 units (*i.e.,* 85%) each time there are not enough REs left in the dataset after the filter (*e.g.,* n of REs < *k*). We ran this sampling program up to n = 1000 times for a *k* range of 1 to 10, and calculated the percentage of cells for which at least one exclusive combination for the target cell type was found. This percentage served as an indicative of the cellular specificity of combinations of *k* active REs.

### Random sampling of intersecting active REs

Using promoter data from the subpanel of 154 primary human cell types, we find that cellular specificity of *k* intersecting REs increases logarithmically from 10-20% for *k* = 1 up to a plateau of 40-80% starting at *k* = 5, depending on the activity threshold (0.5-2 TPM, with a fixed inactivity threshold at 0 TPM; Figure 3B). The 0.5 TPM activity threshold gave the highest selectivity. Relaxing the inactivity thresholds from 0 to 0.1 TPM (with a fixed activity threshold at 1 TPM) increased the % of cells that could be detected by 10-15% depending on the *k* used, again reaching a plateau at around *k = 5* (Figure 3C). A similar scenario was observed using enhancer data, albeit the activity threshold that gave the highest selectivity was lower (0.1 TPM) than for promoters, likely reflecting the generally lower TPM values of the enhancer subset (Figure 3D). Relaxing the inactivity thresholds up to 0.1 (with a fixed activity threshold of 0.5 TPM) did not improve the cell selectivity (Figure 3E). These results suggest that combinations of just a handful of active REs could provide substantial cellular resolution in human. As predicted, the usage of a single input (k = 1) has a very limited potential to detect cell types or cell states. Moreover, even though a two-input AND gate greatly increases the number of detectable cell types, it is unlikely to provide the breadth required to be applicable for a technique aimed at detecting most cell types and/or states in the human body. Finally, at least for this dataset and methodology used, our results suggest that our ability to sort cell types based on active RE intersections plateaus between 4-6 REs.

Safety is also a concern when considering possible human applications of RE activity-based methods, such as unwanted leakage (noisy or unpredicted RE activity) in cell-targeted therapies. Using high *k* values would be beneficial in this sense, because, for each extra *k*, there is an extra safety layer to account for false negatives when compared to *k* = 1. Namely, the probability *p* of leakage decreases exponentially by *p^k^*. By applying the simple RE selection criteria described above (with activity thresholds of 0.5 and 0.1 TPM for promoters and enhancers and a strict inactivity threshold of 0 TPM for both), the usage of a four-input AND gate (*k* = 4 combination of promoters and/or enhancers), which can theoretically add as many as three safety layers against false negatives when compared to *k* = 1, is able to discern ~77% and ~76% of human cell types, respectively (Figure 3B and 3D), suggesting that it is a good compromise between technical feasibility (i.e., generating biological systems that use four REs and translate the activity of the gene products regulated by these REs into a single genetic readout) and breadth of cell types that can be detected. These multiple-input AND gates can also be seen as the minimal intersection of co-activated REs that is diagnostic of a given cell type or state within a given complex mixture of cells in a culture dish, in a tissue biopsy sample, or in the human body. We call these intersecting active REs, <u>V</u>ersatile <u>En</u>try <u>Codes</u> (VEnCodes) (Figure 3F).

### Heuristic selection of intersecting active REs (VEnCodes)

The random “sampling” method still falls short of probing the enormous landscape of possible VEnCodes. We thus attempted a heuristic approach to probe the VEnCode landscape. We used the same binarization criteria as in the sampling method but removed the filter for sparsely-active REs that retains REs that were active in a percentage of the cell types assayed. This was done since the effectiveness of this approach is not affected by a large dataset of less sparsely-active REs. REs occupy the rows of the database and can be represented as 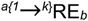 where “*a*” represents the position of the RE in the VEnCode (e.g., for a VEnCode with *k* intersections, *a* will go from 1 to *k*) and *b* represents the row number in the RE list. We then applied a greedy algorithm that considers the sparseness of expression (Figure 4A). In brief, the REs are first sorted by expression sparseness and the sparsest RE (RE1) is chosen as a first-order position (hereafter, “node”) ^1^RE_1_. All cell type columns in which ^1^RE_1_ activity is 0 are then culled from the database, and all remaining ^>1^RE_>1_ are resorted in ascending fashion according to the number of cell types they share co-activity with ^1^RE_1_. Then, ^1^RE_1_ is tested in combination with the next RE (^2^RE_2_) to verify if it satisfies criteria as a VEnCode (*i.e.*, if the intersection between the active REs ^1^RE_1_ ∩ ^2^RE_2_ occurs exclusively in the target cell samples). It follows that for each *k* = 2 combination that satisfies VEnCode criteria, all further *k* > 2 combinations that use these two REs will satisfy the criteria for VEnCode. If no *k* = 2 combination satisfies VEnCode criteria, the algorithm creates secondary nodes and reiterates the pattern described above. To increase the coverage of the landscape, each multiple node test is performed with the three nearest neighbors by order of sparseness. If no *k* = 3 combination satisfies VEnCode criteria, the algorithm creates tertiary nodes and so on. We call this approach the “heuristic approach”.

**Figure 4.**
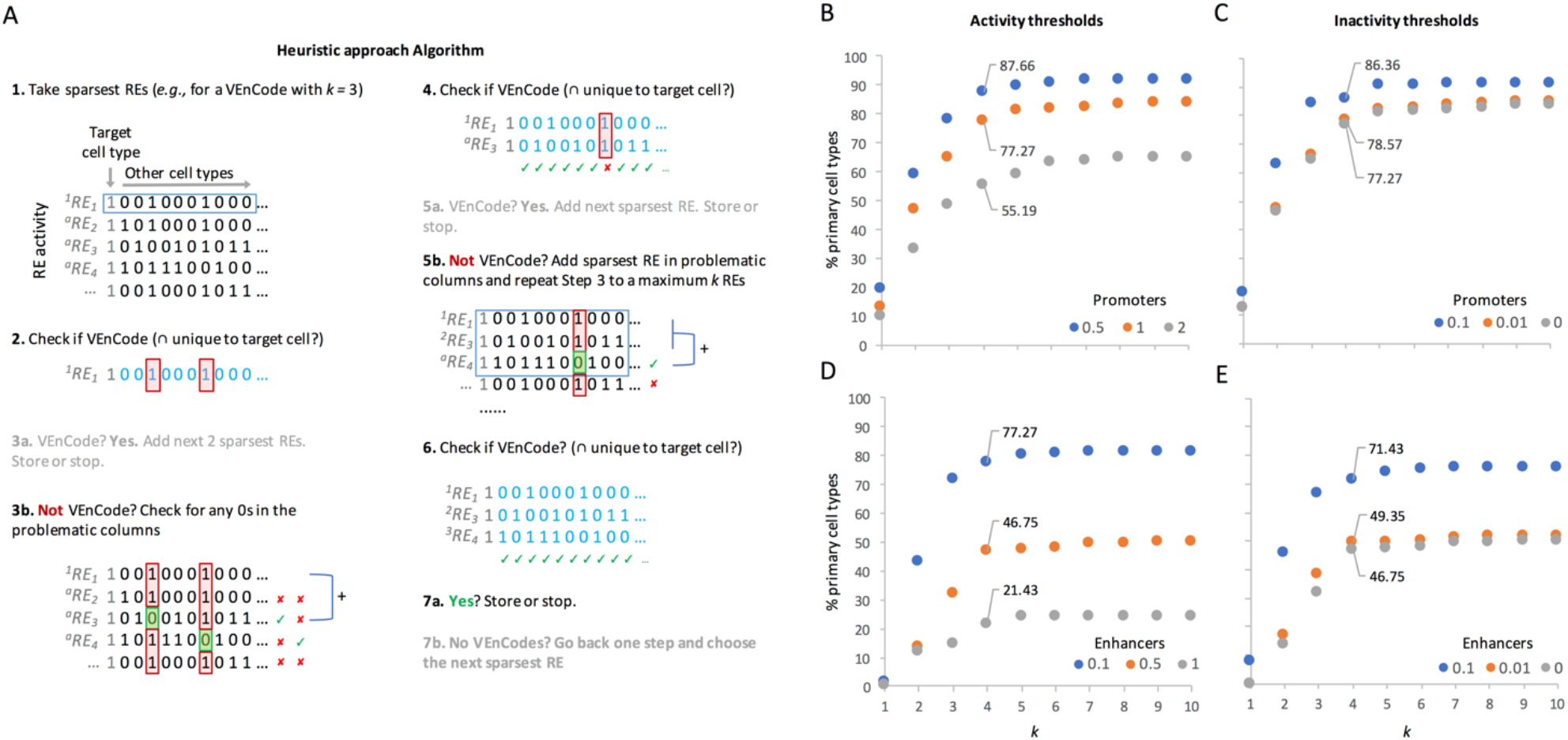
Heuristic method to find intersecting active REs (VEnCodes). **A.** Rationale for the heuristic method. An example is given for a VEnCode with *k* = 3. This algorithm follows a greedy strategy where at each node of the decision tree it makes the locally optimal choice. First, it sorts the REs in the dataset by sparseness, then it takes the sparsest RE (first-level node) and asks if it is inactive in all non-target cell types. If yes, this RE is cell-type specific, and the next *k*-1 sparsest REs can be added to increase safety. If not, it finds out in which cell types this RE is active and searches the data set for a new RE that is inactive in those problematic cell types. If this is successful, then the intersection between these two REs will be specific for the target cell type. In case there is no RE that matches the query, it re-orders the REs by sparseness, this time calculating sparseness only at the “problematic” cell types. It then chooses the sparsest RE as the second-level node and repeats the procedure as described for the first node, increasing node depth until a VEnCode is found. Node depth is always ≤*k* and the algorithm tests several nodes at each level before it gives up. In the example given, there was no need to reorder by sparseness as there was a satisfactory VEnCode. **B-E.** Probing the intersection genetics landscape for promoter **(B, C)** and enhancer **(D, E)** datasets using the heuristic method. Plotted are the percentages of cell types found to have at least one VEnCode per *k* and different activity **(B,D)** and inactivity **(C,E)** TPM thresholds. For the activity panels, the inactivity threshold was fixed at 0 for both promoters and enhancers. For the inactivity panels, the activity thresholds were fixed at 0.5 and 0.1 for promoters and enhancers, respectively.

Applying the heuristic approach to search for VEnCodes using similar threshold conditions as used for the sampling method above, we obtained cell-specific combinations of *k* promoters and enhancers for ~90% and ~80% of the cell types, respectively (Figure 4B-E). More importantly, this method shifts leftwards the plateau for the maximum number of cell types detected so that we are now able to retrieve specific combinations for a larger percentage of cell types even at lower *k* numbers. For instance, at *k* = 4, we retrieve ~88% and ~77% of cell types, using promoters and enhancers, respectively.

To try to find VEnCodes for the cell types where they could not be retrieved using either the sampling or the heuristic method, we combined enhancer (*k_1_*) and promoter (*k*_2_) data in a method we called “heuristic2” approach (Figure 5A). This method increases cellular resolution to ~85% of cell types for *k = 2* (combinations of 2*k_1_* enhancers and 2 *k_2_* promoters) and >90% of cell types for *k* = 4 (combinations of 4 *k_1_* enhancers and 4 *k_2_* promoters, Figure 5B), allowing the generation of VEnCodes for difficult cell types that could not be resolved using promoter or enhancer data alone (Figure 5C). Even though none of our methods saturate the RE activity intersection landscape, these results consistently indicate that combinations of just a few active REs could provide substantial cell type resolution in human.

**Figure 5.**
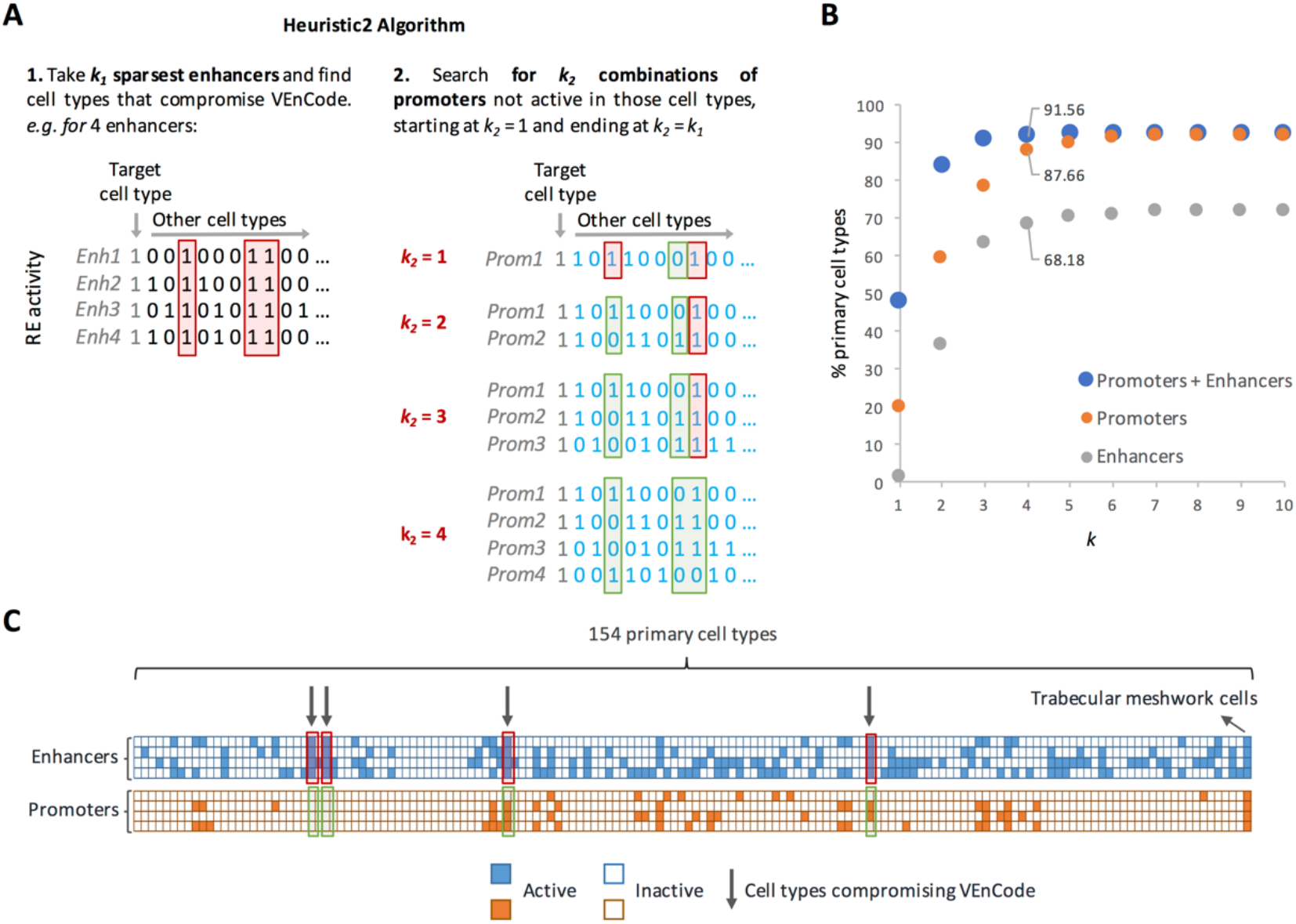
Heuristic2 method to find intersecting active REs (VEnCodes). **A.** Rationale for the Heuristic2 method. This algorithm combines the efficiency of the heuristic method with the extra flexibility of using both enhancers and promoters to target a cell type. First, it finds the *k* sparsest enhancers (*k_1_*) that are active for the target cell type and asks if they are a VEnCode. If they are not, it focuses on the “problematic” cell types in which the enhancers are active, and, using the approach described in Figure 4, asks if there are any combination of promoters (*k_2_*) that are not active in those cell types. If so, then the intersection of the enhancer and promoter activities is specific to the target cell type. **B.** Probing the intersection genetics landscape using the Heristic2 method. Plotted are the percentages of cell types found to have at least one VEnCode per *k* using promoter (orange circles), enhancer (gray circles), or promoter + enhancer (blue circles) data. **C.** Visual representation of a VEnCode obtained using the Heuristic2 method for trabecular meshwork cells. Binary heatmap where each column represents one of the 154 primary human cell types and each row one RE from the enhancer (blue boxes) and promoter (orange boxes) datasets. Red boxes and arrows depict the cell type data that are preventing the interception of enhancers from being a VEnCode for the trabecular meshwork cells. Green boxes highlight the promoter expression data in those problematic cell types.

### Measuring VEnCode robustness

Next, we asked whether we could devise algorithms to rank a VEnCode according to its quality and robustness. A *k =* 4 VEnCode assumes, based on the available RE usage data, that the four chosen REs are never active together in any cell type and/or state except in the desired target cell type and/or state. Clearly, there could be many instances when this premise is false, so that the VEnCode falls apart. For instance, the VEnCode is compromised if the VEnCode is also able to detect a cell type which is not included in the database used or if false negatives are a prevalent artifact of the databases used to devise VEnCodes (*e.g.*, a given RE is labelled as inactive in our database, but it is, in reality, active or for any reason unstably fluctuates between active and inactive states). To attempt to quantify these problems, we carried out Monte Carlo simulations of false negative results by randomly activating REs and recalculating whether or not the VEnCode continued being selective for our target cell type after each simulation (Figure 6A). We scored how many false negatives on average (for *n* simulations) are required until the VEnCode falls apart. This gives the quality value *E_raw_* for each VEnCode. *E_raw_* varies as a function of *k* comprising the VEnCode and the number of cell types *c* in the database. The higher the *k*, the higher *E_raw_*, attesting to the fact that intersections are more robust to technical errors and biological noise. To make *E* comparable between different conditions, we normalize *E_raw_* according to a reference best-case-scenario *E_best(c, k)_* value, which was obtained by Monte Carlo simulations performed as described above, yet for the best-case-scenario for a VEnCode: where all *k* REs are inactive in the nontarget cell types). Hence, normalized *E* = 100\**E_raw_/E_best(c, k)_* (Figure S2 and Table S2). The idea is that *E* is directly proportional to the intraindividual robustness of a given VEnCode towards a cell type (Figure 6B).

**Figure 6.**
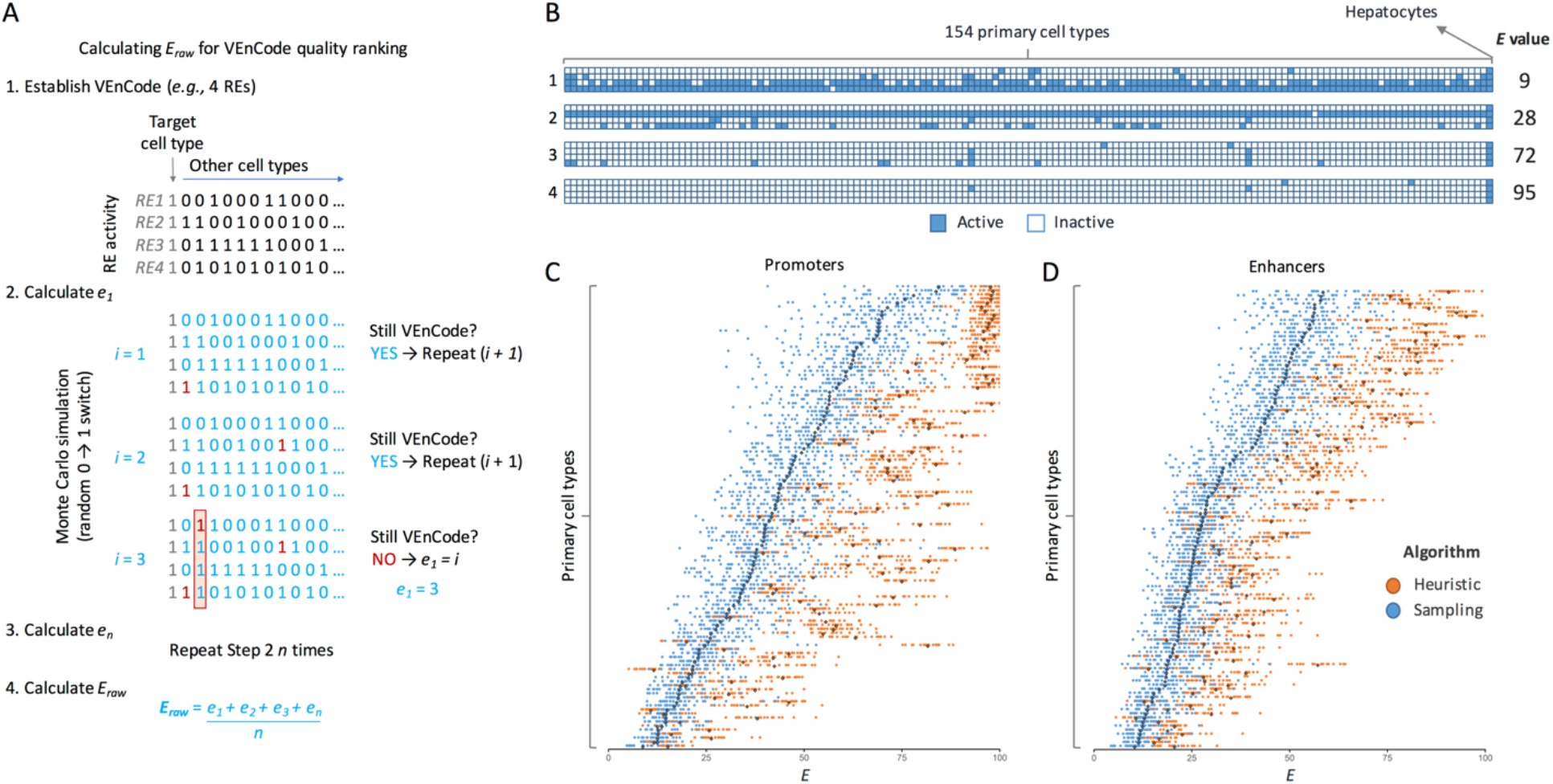
Method for ranking VEnCode intraindividual robustness. **A.** Outline of the method to calculate the *E* value of a VEnCode. *E_raw_* is calculated by taking a VEnCode (1.) and accounting for possible false-negatives in the data by turning inactive REs into active ones (2.). To this end, the algorithm performs random 0-to-1 changes in the dataset, one at a time, and then checks if the VEnCode condition is still satisfied. It reiterates *e_1_* times until the VEnCode condition is no longer satisfied. It then repeats the simulation *n* times (3.) and returns *E_raw_* by calculating the average of all *e* values obtained (4.). *E_raw_* is then normalized according to the formula described in Figure S2 and Table S2 to obtain *E.* **B.** Visual representation of four (1-4) hepatocyte VEnCodes obtained using different algorithms and promoter data. Binary heatmap where each column represents one of the 154 primary human cell types and each row one RE from the promoter data (blue boxes). The *E* value of each VEnCode is depicted on the right. **C-D.** The effect on *E* values of using sampling (blue) or heuristic (orange) methods to obtain VEnCodes. The heuristic method increases average *E* for most cell types for promoter (C) and enhancer (D) data. *y* axis represents different cell types ordered by increasing *E* obtained by the sampling method. Each dot is a VEnCode (n = 5-20 per primary cell type). Darker diamonds represent the mean.

To understand how *E* scales with cell type identity, we used the sampling method to obtain an unbiased set of VEnCodes using *k* = 4 promoters. From the 114/154 cell types for which we retrieved 5-20 VEnCodes in *n* = 10000 samplings, we obtained *E* values varying between 6 and 99 (Figure 6C and Figure S3A). The *E* quality index varied substantially between cell types. For instance, “Fibroblast - Mammary” cells only allow the generation of VEnCodes with small *E* values (between 5 and 17), while hepatocytes allow the generation of high-quality VEnCodes with large *E* values (between 62 and 91). To test whether the heuristic method improved VEnCode quality, we calculated *E* from a subset of 5-20 promoter VEnCodes obtained from 131/154 cell types, which comprised 113 cell types for which we obtained VEnCodes using the sampling method (Figure 6C and Figure S3B). As expected, the heuristic method statistically significantly improved VEnCode quality by an average of 21.1 units (range 6-57) above random sampling for 88.5% of cell types (100/113, *p* < 0. 0005, Bonferroni-corrected unpaired *T* tests, Figure 6C). Similar results were obtained for enhancer VEnCodes: average improvement of 14.1 units (range 4-40) over random sampling for 83% of cell types (93/112, *p* < 0.00005, Bonferroni-corrected unpaired *T* tests, Figure 6D and Figure S3C-D). We conclude that the heuristic method not only finds VEnCodes for a larger amount of cell types, but also generates higher quality VEnCodes.

### VEnCode interindividual robustness

An ideal VEnCode retains its specificity towards the target cell type across multiple individuals of a population. For this, the VEnCode must be robust to interindividual variation on cell-specific RE usage patterns. Interindividual variation could arise either due to technical variation introduced during determination of active and inactive REs for a given cell type in a given individual or as true biological variation in RE usage for that cell type between individuals. The likelihood of relying on false positive calls to generate VEnCodes should be inversely proportional to the number of individuals surveyed for RE usage in the target cell type. To verify this, we estimated interindividual robustness *“z”* of VEnCodes by calculating the percentage of VEnCodes generated from a subset of cell type donors that retained VEnCode satisfiability for all other donors of that cell type, whose data were not used to generate the initial VEnCodes (Figure 7A). Our results show that despite some variability between interindividual robustness across different cell types, on average, their VEnCodes (k = 4) are robust (Figure 7B). Namely, when promoter usage data from one and two donors are used, the *z* values increases on average by ~9.4% from 90.6 to 100%, respectively (*p* < 0.00001, Wilcoxon test for a subset of 66 cell types with 3 donors with 0 < n < 71 VEnCodes generated by the sampling method in all conditions; Figure 7, top left panel). Similar results were found for a subset of cell types (n = 9) with 4 donors, where 2 donors were sufficient to saturate *z* (Figure 7, bottom left panel). Enhancer data from a single donor seems to carry even more predictability for other donors than promoter data, as *z* is on average only 1.4% lower than 100% when data from one donor is used instead of 2 (*p* = 0.00096, Wilcoxon test for a subset of 67 cell types with 3 donors Figure 7, top right panel). When the subset of 6 cell types with enhancer data from 4 donors was analyzed, data from a single donor was sufficient to saturate *z* (Figure 7, bottom right panel). These results suggest that using data from more than one donor is most helpful for promoter data, where it can help significantly increase VEnCode interindividual robustness, and hence the likelihood that a VEnCode will be specific for the target cell in different individuals.

**Figure 7.**
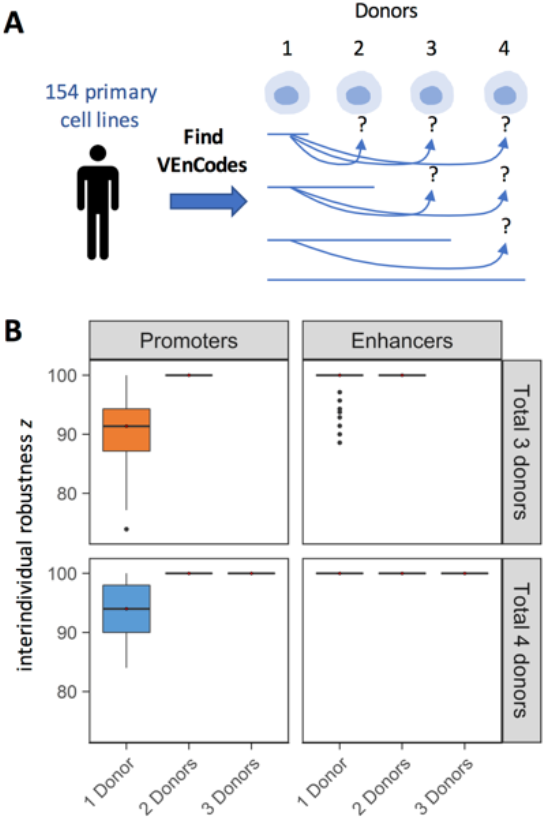
VEnCode interindividual robustness. **A.** Rationale for the estimation of VEnCode interindividual robustness. VEnCodes that are generated based on data from one or more donors are tested as VEnCodes on data from other donors. The % of VEnCodes that satisfy VEnCode criteria for the other donors is *z*. **B.** Box plots representing *z* obtained from various primary cell types based on promoter and enhancer data. Subsets of primary cells for which three (top panels, orange) and four (bottom panels, blue) donors were tested. *z* is saturated at 100% for all cell types tested when VEnCodes are determined using data from 2 cell donors.

Even though there is no correlation between average VEnCode quality *E* for a cell type and the cell type’s interindividual robustness *z* (Figure S4), consistent with the fact that an interindividually robust VEnCode needs not be of high *E* quality, or that a VEnCode with a high *E* score is not necessarily the best VEnCode for multiple individuals, the optimal scenario would be to determine VEnCodes from a large cohort of donors of a cell type and then choose the VEnCodes with highest *E* scores from this subset. With this in mind, we calculated the five best VEnCodes using the Heuristic2 method with for *k* ranging from 1 to 4 for a list of primary cell types with at least 3 donors (Supplementary Data S1). This list can serve as a starting point to explore other properties of VEnCodes and to perform crossvalidation experiments using independent techniques, similar to what we report further below.

### VEnCodes for alternative cell states: cancer

The FANTOM5 database contains RE usage data for 274 cancer cell line samples (Andersson *et al.*, 2014 and FANTOM5 Consortium *et al.*, 2014; Lizio *et al.*, 2015), which can be merged into 158 cancer cell types (Table S3). If VEnCodes could be determined for cancer cell types, they could be used in different cell targeting methods, such as to improve cell selectivity in gene therapy directed towards cancer cells. To verify if VEnCodes could be determined for cancer cell types, we created *in silico* models for diseased patients carrying one cancer cell type each by adding the cancer cell type data to the 154 primary cell type database (Figure 8A). While there are many caveats and sources of additional noise with this strategy, such as cell line heterogeneity, long-term cell culture artifacts, cell donor gender, and the incompleteness of the cell type and state database used, to cite just a few, it already serves the purpose of submitting the cancer cell types to the same stringent criteria as if they were a new primary cell type. Furthermore, the availability of data from cancer cell types obtained from multiple donors provides the possibility to test for interindividual robustness of the cancer cell VEnCodes.

**Figure 8.**
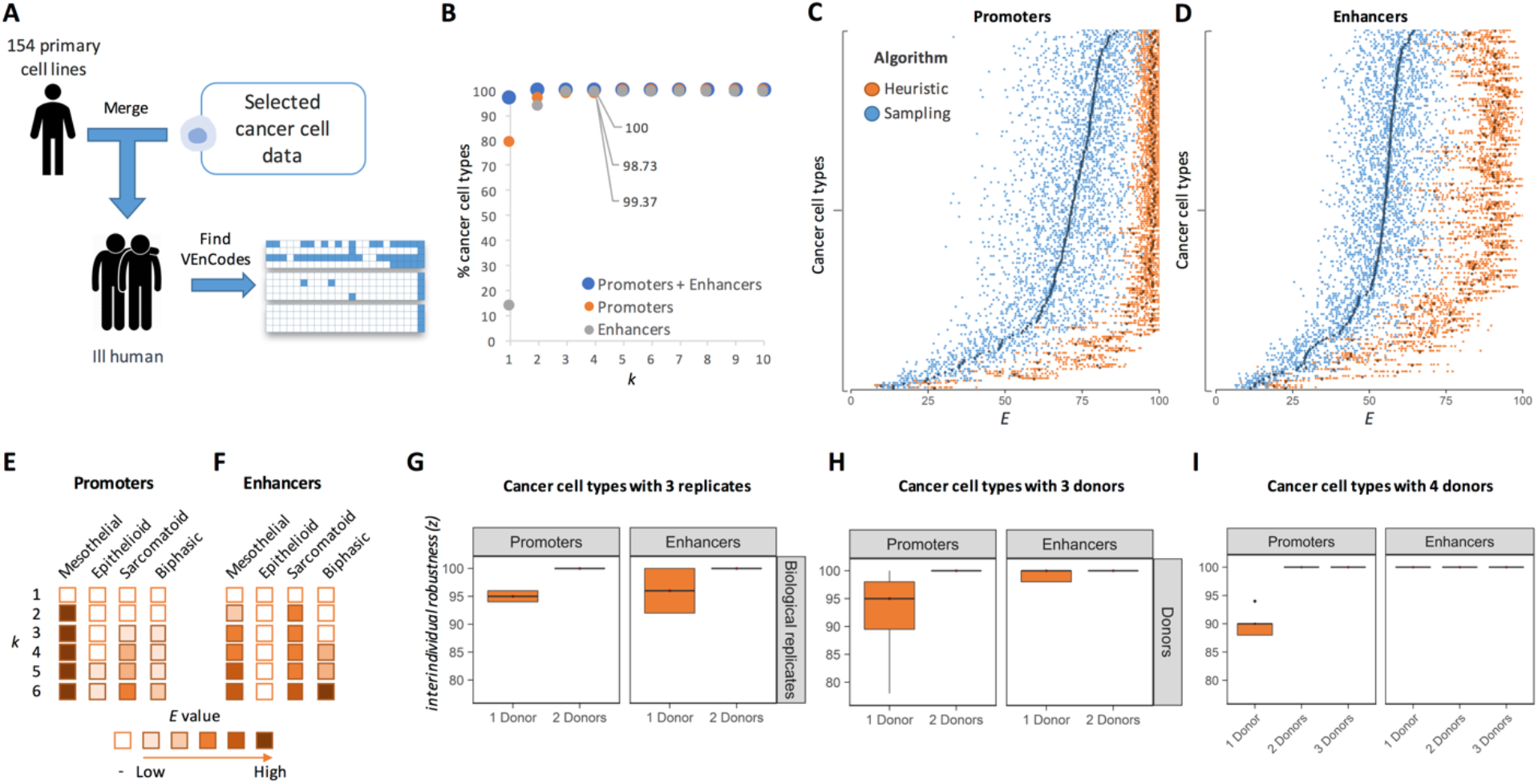
VEnCodes for cancer cell types. **A.** Strategy for simulating a cancer patient in silico. **B.** Probing the intersection genetics landscape for cancer cell types using the Heuristic2 method. Plotted are the percentages of cell types found to have at least one VEnCode per *k* using promoter (orange circles), enhancer (gray circles), or promoter + enhancer (blue circles) data. **C-D.** The effect on *E* values of using sampling (blue) or heuristic (orange) methods to obtain VEnCodes for cancer cell types. The heuristic method increases average *E* for most cell types for promoter **(C)** and enhancer **(D)** data. *y* axis represents different cancer cell types ordered by increasing *E* obtained by the sampling method. Each dot is a VEnCode (n = 5-20 per primary cell type). Darker diamonds represent the mean. **E-F.** Case study of mesothelioma cancer cells stratified into epithelioid, sarcomatoid, and biphasic subtypes. Primary mesothelial cells are shown in the left column as a reference. Rows depict increasing *k.* Boxes are filled if at least one VEnCode is found using *k* REs. If a VEnCode is found, the box is colored according to binned average *E* value of the VEnCodes found (n = 1-20). **G-H.** Box plots representing interindividual robustness *z* values obtained from all cancer cell types with 3 **(G, H)** or 4 **(I)** donors based on promoter (left panels) and enhancer (right panels) data. **G.** Subsets of cancer cells for which biological replicates were available (*i.e.*, repeated assays with the same cancer cell line). **H-I.** Subsets of cancer cells types for which independent cell lines were analyzed. *z* is saturated at 100% for all cell types tested when VEnCodes are determined using data from 2 cell donors.

Exploring the RE landscape of cancer cell types we again noticed that VEnCodes are readily obtained even for smaller *k* values (Figure 8B), except for enhancers, where only ~14% of cancer cell types had a specific enhancer (*k* = 1, Figure 8B). At *k* = 4, ~99% of cancer cell types surveyed could be distinguished using the heuristic method for promoters or enhancers (Figure 8B). This goes up to 100% using the heuristic2 method already with a *k* = 3 (Figure 8B). Cancer cell type VEnCodes are generally of very high quality, as shown by their large *E* values (Figure 8C and 8D). Using the heuristic method increases the *E* values, similarly to what we observed in primary cell lines (Figure 8C and 8D).

One caveat of the *in silico* cancer patient model is that not all cells of origin of some cancer cell types are present in the primary cell database. This is the case for small cell lung carcinoma (SCLC), which is thought to originate from neuroendocrine cells of the lung (Park *et al.*, 2011). Certainly, an expansion of the primary cell database is warranted and it would help generate safer and more robust VEnCodes.

To study this issue more carefully, we looked at mesothelioma, for which the assumed primary cell of origin, the mesothelial cell, is available in the current database. We first stratified the mesothelioma cell types into three cytological classes according to Cellosaurus (Barioch, 2018): epithelioid (n = 7: ACC-MESO-1, ACC-MESO-4, Mero-14, Mero-41, Mero-82, Mero-95, NCI-H226, and No36 (epithelial-like stellate cells)), sarcomatoid (n = 3: NCI-H2052, NCI-H28, and ONE58), and biphasic (n = 5: Mero-25, Mero-48a, Mero-83, Mero-84, NCI-H2452). We then asked how difficult it was to generate robust VEnCodes for these mesothelioma types (Figure 8E and 8F). We find that while VEnCodes can be readily generated for primary mesothelial cells with *k* = 2, larger *k* values are required to generate VEnCodes for mesothelioma cells. VEnCodes were found for all mesothelioma subtypes, except for epithelioid mesothelioma cells, which could only be identified when promoter data was used, and even then they were of poor quality (E = ~7). In general, VEnCode intraindividual robustness *E* increased with higher *k*, again attesting for the potential safety value of using more intersections (Figure 8E and 8F).

As many cancer cell types are characterized by a level of heterogeneity, we were expecting less interindividual robustness in cancer cells relative to primary cell types. We thus applied the sampling method to calculate the interindividual robustness *z* of cancer cell types. We found that cancer cell type VEnCodes (*k* = 4) determined either from promoters or enhancer usage data, have very high interindividual robustness *z*, which is already saturated when data from 2 donors are used (Figure 8G-I). These results show that small RE usage signatures can reproducibly define dozens of cancer cell types. The level of interindividual robustness is similar to that of technical replicates (Figure 8G, compare top and bottom panels). Even genetically hypervariable cancer cell types, such as SCLC cells (George *et al.*, 2015), for which data from four cell lines were available, also gave 100% *z* values when data from two donors were used (Figure 8I). We conclude that highly robust and safe cancer cell VEnCodes can be obtained using CAGE-seq data.

### VEnCode cross-validation

Having shown that a publicly-available RE usage database based on CAGE-seq data can be used to generate and quality-rank VEnCodes for hundreds of primary and cancer cell types, we next asked whether these CAGE-seq-based VEnCodes could be cross-validated using other publicly-available comprehensive RE usage datasets. There are two types of cross-validation that would be desirable: first, to show that all the REs used in the VEnCodes for a given cell type are indeed active in that cell type. Second, that the combination of enhancers is exclusively active in that cell type. Of these, only the first type of cross-validation is currently feasible without extensive consortium-level biological experimentation due to the lack of a suitable database with the breadth and depth of the FANTOM5 database as regards RE usage in primary cells. To cross-validate CAGE-seq VEnCodes for a specific cell type using RE activity estimated by other methods in the same cell type, we searched the literature for suitable studies and found 18 candidate cell types that could be used for cross-validation (Table S4): three “healthy” cell types (human induced pluripotent stem cells (hiPSCs) and 2 primary cell types), and 15 cancer cell types/lines. Whereas FANTOM5 data on hiPSCs were not included in our curated primary cell database, the fact that hiPSCs and human embryonic stem cells (hESCs) share nearly identical molecular profiles and pluripotency properties (Chin *et al.*, 2010; Bock *et al.*, 2011; Marei *et al.*, 2017) allows the usage of a high-quality functional enhancer dataset generated for hESCs (Barakat *et al.*, 2018) for VEnCode crossvalidation. The methods for RE activity estimation in the retrieved studies varied between chromatin immunoprecipitation (ChIP)-based methods, DNA accessibility-based methods (FAIRE, DHS, ATAC-seq), enhancer function methods (eRNA, STARR-seq), and combinations of these methods (Table S4). We downloaded raw data (BED, FASTA, CSV, BroadPeak, or TSV files) from the studies and parsed the data to retrieve compatible genomic locations of “active” or potentially active enhancers (for DNAse accessibility-dependent techniques).

To cross-validate the CAGE-seq-determined VEnCodes, we generated up to 200 VEnCodes (with *k* = 4) for each of the 18 cell types and determined the fraction of *k* per VEnCodes that were considered active in the external database (Figure 9A). Any degree of overlap (>0 nt) between the CAGE-seq RE coordinates and the external database active RE coordinates was considered positive for validation. Cross-validation results varied between cell types and studies. Fully validated *k* = 4 VEnCodes, for instance, were found for 11/18 (61.1%) cells, while partially-validated *k* ≥ 2/4 in 17/18 (94.4%) cell types (Figure 9B). The low validation for some cell types could be due to many factors, such as cell identity and the nature of the method used to determine RE activity. Indeed, the likelihood of VEnCode cross-validation correlated significantly with the % overlap of active RE calls between CAGE-seq and the external method (r = 0.90, *p* < 0.00001; Figure S5). This is not unexpected as the readouts used to determine enhancer activity in the different studies vary in specificity and sensitivity. Regardless of these limitations, our results indicate that for a majority of cell types, CAGE-seq VEnCodes can be cross-validated using RE usage datasets independently determined by other methods. To bypass the limitation of low overlap between enhancer activity calls in different database, we first cross-validated enhancers, and then determined VEnCodes using the curated primary cell CAGE-seq data (Figure 9C). With this approach, cross-validated VEnCodes were obtained for all 18 cell types (Figure 9D), and these VEnCodes could be further quality-ranked according to their E value (Figure S6 and Supplementary Data S2). Hence, high-quality VEnCodes can be found using exclusively the subset of CAGE-seq REs that are cross-validated against publicly-available RE usage datasets.

**Figure 9.**
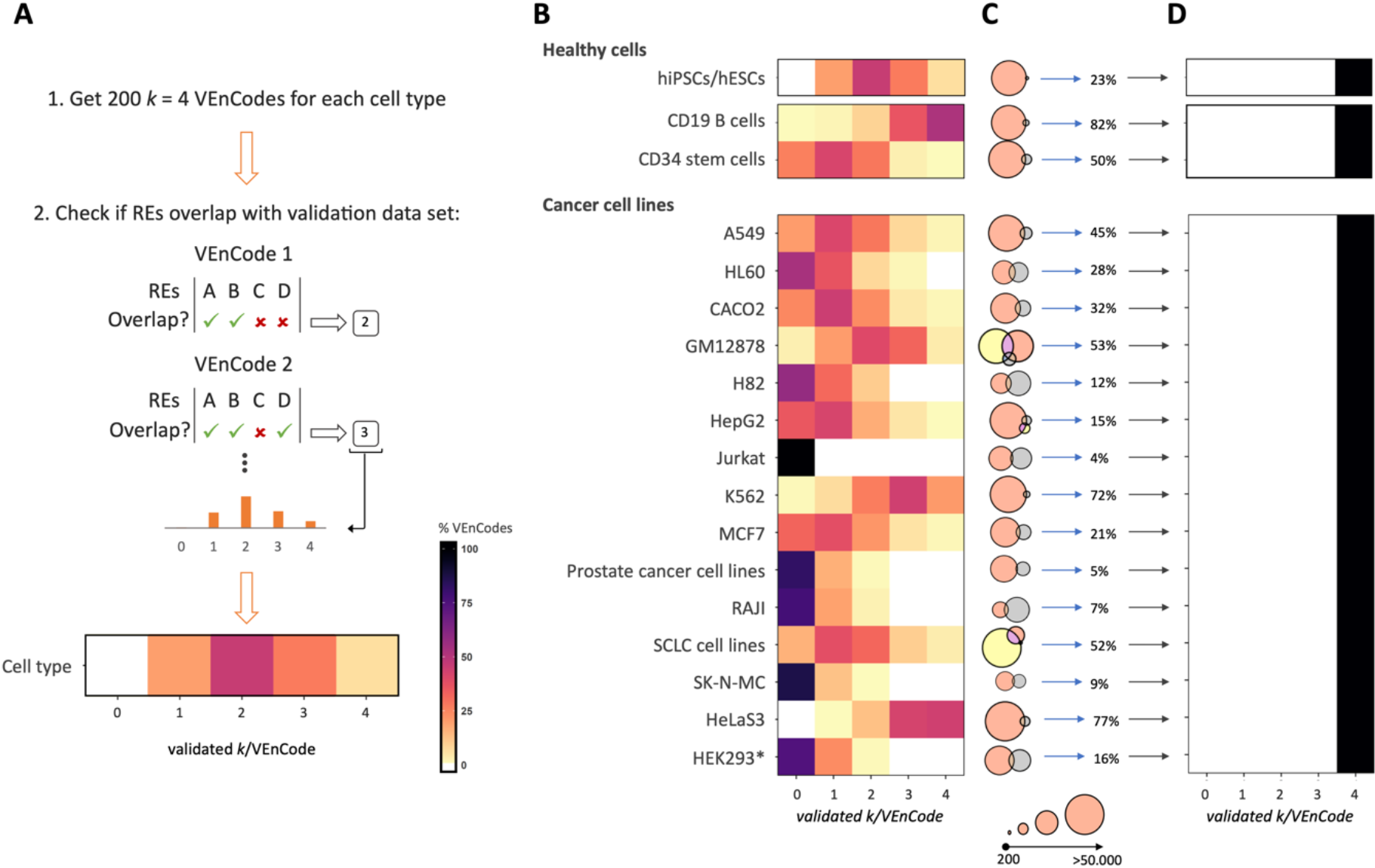
Cross-validation of CAGE-seq-determined VEnCodes. **A.** Strategy for cross-validation of CAGE-seq-determined VEnCodes using external databases. **B.** Heatmaps depicting the distribution of cross-validated VEnCodes, for *k* = 4 enhancers, according to the number of validated *k*/VEnCode for each cell type. The databases used for cross-validation of each cell type and the full description of the cell types are provided in Table S4. **C.** Venn diagrams depicting the % of active enhancer calls determined in this study using the FANTOM5 database CAGE-seq data (gray circles; notice that they are sometimes too small to see in the figure) that overlap with the external cross-validation databases (peach and yellow). **D.** Heatmaps depicting the distribution of cross-validated VEnCodes, for *k* = 4 enhancers, according to the number of validated *k*/VEnCode for each cell type using exclusively CAGE-seq enhancers that are present in the external databases. *HEK293 cells originate from fetal kidney tissue, but they were placed in this group as they are derived from adenovirus-transformation and have a complex karyotype.

### VEnCodes using single-cell sequencing data

Genomewide RE activity profiles have traditionally been obtained using large cell populations, as in the case of the FANTOM5 CAGE-seq data studied herein. While the advantage of these bulk preparations is the increased depth and resolution of the RE activity predictions obtained, a clear disadvantage is the loss of single-cell resolution. This is most evident in situations where RNA is prepared from complex mixtures of cells, such as healthy tissues samples and cancer biopsies. Single-cell strategies that can both resolve cell heterogeneity and infer RE activity have been developed, but they still provide a relatively shallow (discontinuous) and noisy view of RE activity per cell (e.g., Buenrostro *et al.*, 2015; Cusanovich *et al.*, 2015; Chen *et al.*, 2018; Cusanovich *et al.*, 2018a; Cusanovich *et al.*, 2018b Mezger *et al.*, 2018; Kuono *et al.*, 2019). These properties limit the usefulness of single-cell strategies for VEnCode determination, which requires very stringent RE activity criteria, especially for negative RE activity calls. Single cell CAGE-seq (C1 CAGE), for instance, recovers per cell on average ~15, ~10, ~5, and <5% of bulk CAGE-seq-determined enhancers expressed at 10-100, 5-105, 1-5 and <1 TPM (Kuono *et al.*, 2019). Considering that the thresholds for enhancer inactivity and activity in our study are 0 and >0.5 TPM, this suggests that the rate of false positives in C1 CAGE-determined VEnCodes would be high.

Some of the limitations described above can be partially circumvented using strategies that consolidate single cell RE activity profiles by pooling a sufficiently large number of single cells after taxonomic cell clustering (*e.g.*, Pott and Lieb, 2015; Alessandri *et al.*, 2019; Bravo-González-Blas *et al.*, 2019; Lareau *et al.*, 2019; Stuart *et al.*, 2019; Urrutia *et al.*, 2019; Xiong *et al.*, 2019; Zeng *et al.*, 2019; Zhou *et al.*, 2019). We thus hypothesized that pooled C1 CAGE Seq data could retrieve enough enhancers to enter the VEnCode determination pipeline.

To test this, we obtained pooled enhancer activity prediction data from a subset of untreated adenocarcinomic human alveolar basal epithelial cell line [“T0” (untreated) A549 cells, n = 35 cells (Table S5); Kuono *et al.*, 2019). From this set of cells, we retrieved 540 unique enhancers, which were considered “ON” (Table S6). All other non-retrieved enhancers were considered “OFF”. This data was then processed as described above for bulk CAGE-sequenced cancer cells (Figure 10A). Results showed that VEnCodes could be successfully retrieved from enhancer activity profiles obtained from pooled C1 CAGE data (Figure 10B). In general, C1 CAGE-based VEnCodes obtained by the sampling method were 34.1 and 38.3% worse in quality than bulk and bulk-validated CAGE-seq-determined VEnCodes, respectively (35.0 ± 6.7, 53.1 ± 7.5, and 56.7 ± 6.90, average ± SD *E* values for C1 CAGE, bulk, and bulk-validated CAGE-seq enhancers, respectively; *p* < 0.01 for each respective comparison; Tukey HSD post-hoc test, Figure 10B and 10C). A similar pattern was observed for VEnCodes obtained using the heuristic method, albeit VEnCode quality was significantly increased for all estimates, as compared to the sampling method, as expected (*p* < 0.01; Tukey HSD post-hoc test, Figure 10B and 10C). We conclude that RE activity profiles obtained from single-cell sequencing data can be successfully integrated into the VEnCode pipeline when pooled estimates are used. At least for C1 CAGE seq of a small number of A549 cells, these estimates are not optimal as those from bulk-sequencing data due to their proneness to false negative RE calls and the generally reduced quality of the VEnCodes generated.

**Figure 10.**
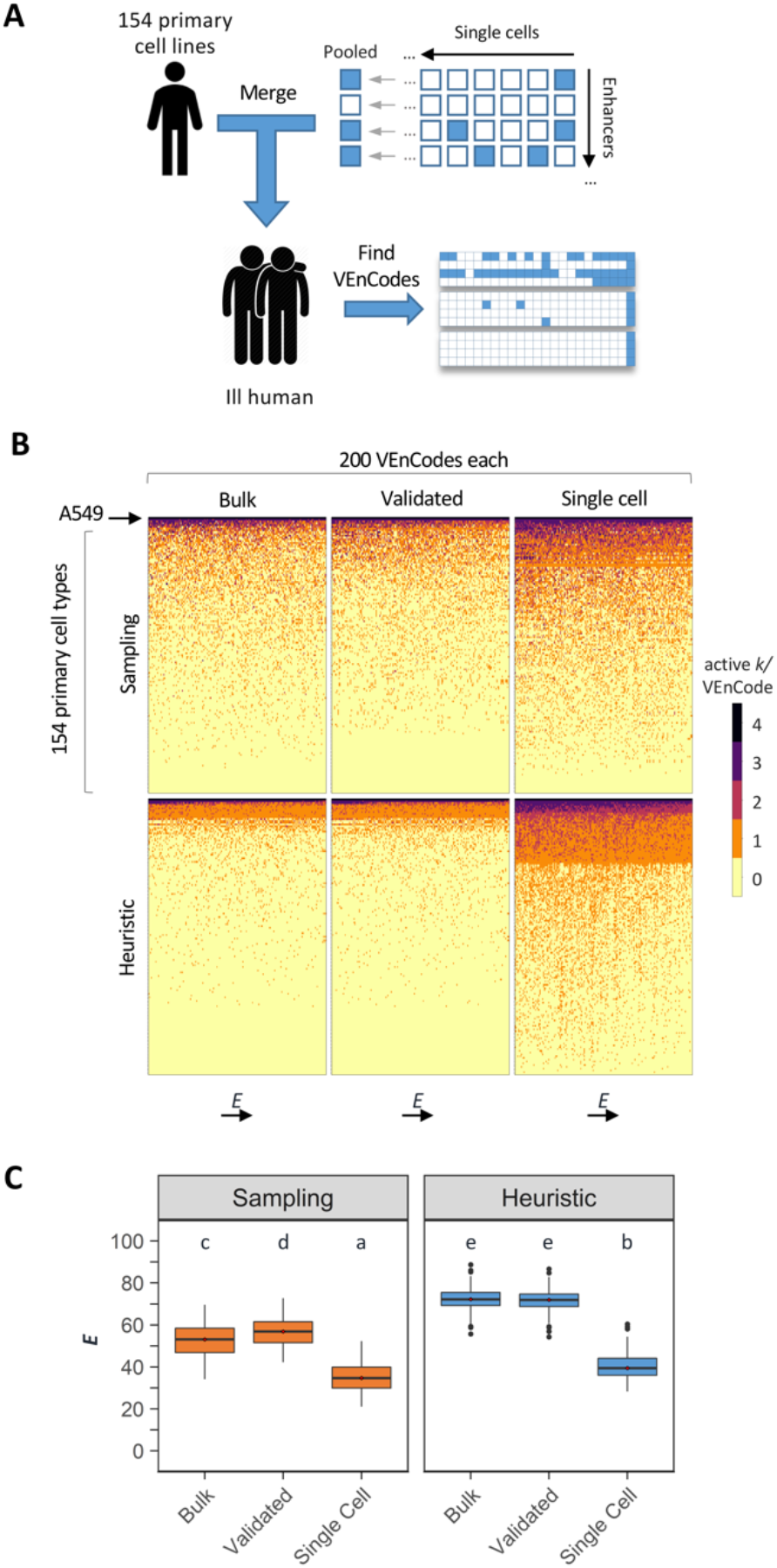
VenCodes from single-cell-derived RE usage data. **A.** Strategy for integrating single-cell-derived RE usage data into the VEnCode-generation pipeline. Single-cell data, such as enhancer usage patterns obtained from C1 CAGE-seq, are pooled and then integrated into the curated database of bulk CAGE-seq enhancer panel. **B.** Heatmaps depicting the *E* quality of 200 *k* = 4 enhancer VEnCodes obtained by the sampling or heuristic method from single cell (35 pooled “T0” cells from C1 CAGE-seq; Kuono *et al.*, 2019), bulk CAGE-seq (FANTOM5), or bulk-validated CAGE-seq data (as described in Figure 9) for A549 cells. Columns represent 200 VEnCodes ranked according to *E* value, and rows represents cell types, ranked according to average ((active *k*)/VEnCode). **C.** Box plots representing the quantification of the *E* value data reported in **B.** Different letters represent conditions that are statistically significantly different (ANOVA performed with all six conditions, followed by Tukey HSD, *p* < 0.01).

## DISCUSSION

A major challenge in biomedicine is to access and gain control of a specific cellular type, be it in a healthy or disease state, within a complex and highly adaptable body. A methodology that allows genetic access to all cellular types and states in the human body would have a major impact in multiple domains of life science, including the possibility of studying and designing novel research tools, therapies, as well as better bioinspired technology and cosmetics. Such methodology addresses a major problem in the fields of life sciences research, biological engineering, and gene therapy: cellular-targeting, i.e., how to restrict the desired genetic intervention to a unique set of cells within an organism or different cell states within unicellular populations.

Even when specific solutions exist (e.g., antibodies against target cell surface proteins or viruses with tropism towards certain cell types) that give access to a single cell type or state in an organism, no approach is known that allows for the systematic generation of similar specific solutions for other cell types or states in any given organism. Therefore, there is a profound limitation in the technologies available to genetically access particular cellular types and states in a very limited set of organisms.

An alternative to these procedures is to use methods that do not rely on cell-specific strategies to deliver genetic materials to cells. For instance, to use a system that delivers the desired genetic material to as many cells as possible in a complex organism, unrestrictedly. Considering that such systems are becoming available (e.g., unbiased non-integrating viral delivery or chemical-based delivery), the challenge becomes to have a highly versatile approach to activate any particular genetic message exclusively within a target cell type.

Intersectional genetics provides a solution for cellular targeting in complex organisms. However, there are several challenges that have to be overcome in order to apply intersectional genetics in human. The first challenge is the understandable lack of a library of gene drivers with known expression patterns to choose intersections from. To overcome this, we attempted to explore alternative resources such as large RE usage databases determined using next-generation sequencing methods. We explored a curated panel of 154 primary human cell types and 158 cancer cell types for which a uniform RE usage atlas consisting of CAGE-seq data is currently available (Andersson *et al.*, 2014; FANTOM Consortium and the RIKEN PMI and CLST (DGT), 2014; Lizio *et al.*, 2015). FANTOM5 data and other datasets have been previously explored as potential sources of cell-specific features, including enhancers (Ienasescu *et al.*, 2016; Gao *et al.*, 2016). One of these tools is SlideBase, which uses interacting sliders for the selection of expressed features from a given dataset by user-customized expression thresholds (Ienasescu *et al.*, 2016). However, while such user-friendly tools can serve this and many other purposes, they are neither conceived nor optimized for a systematic analysis of intersectional genetics. An additional limitation is that the datasets have not been curated with the conservative criteria for unique cell types that we used.

The second challenge is to understand the landscape of intersectional genetics in human, including its safety and reliability. To the best of our knowledge a systematic assessment of the potential and robustness of the intersectional genetics approach had never been performed for any organism, much less for humans. How far could intersectional genetics take us as an approach to gain accessibility to any given human cell? If each cell type does not have a uniquely active RE, or if the usage of a single unique RE carries high risk for therapeutic and diagnostic purposes due to technical artifacts and biological noise, would the unique intersection of two or more REs be enough to generate cell type specific gene drivers for every human cell type? This is a relevant question as there are several technical solutions available to explore genetically the intersections of active REs, such as split-transcription factors and recombinase-based FLP-OUT strategies (Lakso *et al.*, 1992; Struhl and Basler, 1994; Awatramani *et al.*, 2003; Suster *et al.*, 2004; Stockinger *et al.*, 2005; Luan *et al.*, 2006; Farago *et al.*, 2006; Nissim *et al.*, 2007; Nissim and Bar Ziv, 2010; Siuti *et al.*, 2013; Liu *et al.*, 2014; Morel *et al.*, 2016).

We found herein that >90% of the primary human cell types surveyed can be safely and robustly distinguished from each other with as little as 3 to 4 REs. We called these combinations “VEnCodes” (for Versatile Entry Codes). VEnCodes can be defined as the smallest gene expression ON/OFF signature that carries enough diagnostic value to distinguish between the target cell and other non-target cell types within a complex mixture of cells. Clearly, VEnCodes with 1 and 2 REs exist and their technical exploitation is already feasible with current techniques. However, for many cells, more REs are required either to obtain a VEnCode, or to obtain a safer and more robust VEnCode. Hence, new intersectional methods are desirable to capitalize on the intersection of 3 or more active REs.

While we obtained VEnCodes for most cells using heuristic methods, we failed to obtain VEnCodes for ~10% of the primary cells surveyed, even when 10 RE intersections were allowed. It is important to notice that we have by no means saturated the VEnCode search space. Hence, more thorough, brute-force methods (*e.g.*, an exhaustive sampling method) might find VEnCodes for these difficult cell types. However, some of these cell types might indeed have poorly-distinguishable or indistinguishable RE activity profiles. These cell types might require other techniques for detection. One possibility, which was not explored here, is to use other intersectional methods based other Boolean logical operations, such as OR, NOT, NOR.

To create a quality index for VEnCodes we determined its susceptibility to technical artifacts and biological noise using Monte Carlo simulations. We show that average VEnCode quality varies significantly between different primary cell types, so that certain cells, such as mast cells and hepatocytes, are more safely distinguishable from others than most fibroblasts subtypes. By exploring RE usage data from the same primary cell type obtained from multiple donors, we find that VEnCodes are very (~100%) robust, especially when determined using enhancer data. Promoter data-based VEnCodes for primary cell types increases when data from at least two cell type donors are used. It is not clear if this reduced robustness using single donor promoter data is due a technical or biological source of noise.

To probe the RE space in a cell state paradigm, we explored data from different cancer cell lines isolated from patients diagnosed with tumors of the same cellular origin. Several cancer cell types are hypervariable in nature, posing a challenge for finding a specific VEnCode that detects the same cancer cell type across multiple individuals. However, VEnCodes could be determined for all cancer types surveyed where multiple cell lines were available, even for notoriously hypervariable cancer cell types such as SCLC cells. Furthermore, as VEnCode retrieval can be systematized, in the absence of a single VEnCode that satisfies detection criteria for multiple cancer cell subtypes, multiple VEnCodes can be designed to account for cancer cell heterogeneity.

Finally, we showed that VEnCodes obtained from CAGE-seq data can be cross-validated using enhancer usage data obtained by other methods. Intuitively, the methods of choice to cross-validate VEnCodes in target cells are methods that infer enhancer function, such as simple luciferase assays or STARR-Seq, where functional enhancers transcribe themselves (Arnold *et al.*, 2013). Cross-validation of a second key assumption of VEnCodes -that they only function in the desired cell type and not in other cell types-is much more challenging, but can partially be addressed with larger primary cell RE activity databases and panels of complex human organoids.

VEnCodes can now be explored as minimal RE-program-sensing parts that can be encoded genetically into plasmid-based biosensors, packaged into viral or non-viral systems, and delivered to cells in the body to diagnose whether or not the RE program of a given cell matches that of the VEnCode. Engineering biosensors that sense the activity of 3 to 4 REs and then perform a multiple-AND gate computation to generate a single output is technically feasible with synthetic biology. Such genetic biosensors could revolutionize medicine by allowing safe and specific gene delivery to any cell type or cell state in the human body.

Enhancer-based VEnCodes are clearly the most promising combinations for generating intersectional genetics tools. Each enhancer can, for instance, be placed directly upstream of a general or synthetic basal promoter. In contrast, one needs to consider that promoter-based VEnCodes, such as those obtained in the heuristic2 method, might not necessarily autonomously convey the desired cell type specific transcription when placed in a synthetic construct context. Nevertheless, there are many efforts to map enhancer-promoter interactions (*e.g.*, Andersson *et al.*, 2014; Mora *et al.*, 2016; Hait *et al.*, 2018), which could be used to optimize the heuristic2 method.

Finally, we have shown that single cell-based strategies for RE estimation such as C1 CAGE are compatible with the VEnCode pipeline. The VEnCodes obtained were understandably of lower quality than those derived from bulk CAGE-seq data alone. Pooling data from larger numbers of cells can theoretically improve VEnCode quality considering larger numbers of REs are obtained. The low-confidence negative RE activity calls generated by pooled single cell-sequencing data are another point of concern, which might also be mitigated by pooling larger numbers of cells. Nevertheless, a major benefit of single-cell strategies for VEnCode applications, is their potential to significantly increase the number of cell types with RE activity profiles. This potential to molecularly profile cell types and even to discover new cell types has become evident with single-cell RNA-seq (Macosko *et al.*, 2015; Bahar Halpern *et al.*, 2017; Regev *et al.*, 2017; Hon *et al.*, 2018). Expanding the number of cell types in the curated singlecell RE activity database is critical because it helps reduce the amount of false VEnCodes for all cell types (e.g., the larger the amount of cell types with robust known active RE calls, the less likely it is to obtain a VEnCodes that will fail in practice). Hence, the inclusion of a new cell type with a single-cell-derived, partial RE profile based exclusively on sparse active RE calls is better than not having the profile at all. Finally, it would be interesting to verify in future studies whether, apart from integrating single cell and bulk RE activity profiles, the algorithms and strategies described herein could also be applied to RE activity data generated exclusively from single-cell data.

In summary, our results suggest that VEnCodes for a wide-variety of human primary cell types and cancer cells can be discovered, quality-controlled, and cross-validated *in silico* using heuristic algorithms and publicly-available genome-wide RE-usage databases, such as the FANTOM5 promoter and enhancer atlases. VEnCodes could be used to engineer intracellular biosensors or devices that use intersectional genetics tools to “read” the VEnCodes and translate them into a custom genetic output. This would allow systematic genetic access to any of these cell types or states. Genetic access carries enormous therapeutic potential by allowing the selective delivery of genetic messages and cures to cells, such as various forms of gene therapy or the specific genetic ablation of abnormal cancerous cells.

## Author’s contributions

AM and AMG designed the study, implemented analyzes, analyzed the data, and wrote the manuscript.

## Acknowledgements

We thank Drs. Fabiana Heredia, Andres Garelli, Musa Mhlanga, Ney Lemke, Tatiana Torres, and Ms. Benilde Pondeca, and other members of the Integrative Biomedicine Laboratory for important discussions, comments, and/or suggestions on the manuscript. A.M.G. was supported by an FCT Investigator Grant (IF/00022/2012). A.M. was supported by fellowships from FCT (SFRH/BD/94931/2013) and the LIGA PORTUGUESA CONTRA O CANCRO (LPCC) 2017. Work in the Integrative Biomedicine Laboratory is supported by the FCT (UID/Multi/04462/2019; PTDC/BEXBCM/1370/2014; PTDC/MED-NEU/30753/2017; and PTDC/BIA-BID/31071/2017), the MIT Portugal Program (MIT-EXPL/BIO/0097/2017), and FAPESP (2016/09659-3).

## Materials and Methods

Materials (code and data) are available at https://github.com/AndreMacedo88/VEnCode. Python language was used to implement all the algorithms and methods in this study (Python Software Foundation. Python Language Reference, version 3.6.5. Available at http://www.python.org. The R language with the ggplot2 package was used to generate the plots for the Figures (R Core Team, 2013: http://www.R-project.org/; Wickham, 2016).

## Supplementary Materials

Supplementary materials include Supplementary Figures S1-6, Supplementary Tables S1-S6, and Supplementary Data S1-2. The latter are available at https://github.com/AndreMacedo88/VEnCode.

## Conflicts of Interests

The authors declare no conflicts of interests.

## Notes

#### Summary of Updates

We provide methods for the cross-validation of CAGE-seq-derived VEnCodes and for the extraction of VEnCodes from pooled single cell sequencing data.

https://github.com/AndreMacedo88/VEnCode

## References

Alessandrì L, Cordero F, Beccuti M, Arigoni M, Olivero M, Romano G, Rabellino S, Licheri N, De-Libero G, Pace L, Calogero RA. rCASC: reproducible classification analysis of single-cell sequencing data. Gigascience. 2019 Sep 1;8(9). pii: giz105.

Andersson R, Gebhard C, Miguel-Escalada I, Hoof I, Bornholdt J, Boyd M, Chen Y, Zhao X, Schmidl C, Suzuki T, Ntini E, Arner E, Valen E, Li K, Schwarzfischer L, Glatz D, Raithel J, Lilje B, Rapin N, Bagger FO, Jørgensen M, Andersen PR, Bertin N, Rackham O, Burroughs AM, Baillie JK, Ishizu Y, Shimizu Y, Furuhata E, Maeda S, Negishi Y, Mungall CJ, Meehan TF, Lassmann T, Itoh M, Kawaji H, Kondo N, Kawai J, Lennartsson A, Daub CO, Heutink P, Hume DA, Jensen TH, Suzuki H, Hayashizaki Y, Müller F; FANTOM Consortium., Forrest AR, Carninci P, Rehli M, Sandelin A. An atlas of active enhancers across human cell types and tissues. Nature. 2014 Mar 27;507(7493):455–61.

Arnold CD, Gerlach D, Stelzer C, Boryń ŁM, Rath M, Stark A. Genome-wide quantitative enhancer activity maps identified by STARR-seq. Science. 2013 Mar 1;339(6123):1074–7. doi: 10.1126/science.1232542.

Awatramani R, Soriano P, Rodriguez C, Mai JJ, Dymecki SM. Cryptic boundaries in roof plate and choroid plexus identified by intersectional gene activation. Nat Genet. 2003 Sep;35(1):70–5.

Bairoch A. The Cellosaurus, a Cell-Line Knowledge Resource. J Biomol Tech. 2018 Jul;29(2):25–38. doi: 10.7171/jbt.18-2902-002. Epub 2018 May 10.

Bahar-Halpern K, Shenhav R, Matcovitch-Natan O, Tóth B, Lemze D, Golan M, Massasa EE, Baydatch S, Landen S, Moor AE, Brandis A, Giladi A, Stokar-Avihail A, David E, Amit I, Itzkovitz S. Single-cell spatial reconstruction reveals global division of labour in the mammalian liver. Nature. 2017 Feb 16;542(7641):352–356.

Barakat TS, Halbritter F, Zhang M, Rendeiro AF, Perenthaler E, Bock C, Chambers I. Functional Dissection of the Enhancer Repertoire in Human Embryonic Stem Cells. Cell Stem Cell. 2018 Aug 2;23(2):276–288.e8.

Bock C, Kiskinis E, Verstappen G, Gu H, Boulting G, Smith ZD, Ziller M, Croft GF, Amoroso MW, Oakley DH, Gnirke A, Eggan K, Meissner A. Reference Maps of human ES and iPS cell variation enable high-throughput characterization of pluripotent cell lines. Cell. 2011 Feb 4;144(3):439–52.

Bravo González-Blas C, Minnoye L, Papasokrati D, Aibar S, Hulselmans G, Christiaens V, Davie K, Wouters J, Aerts S. cisTopic: cis-regulatory topic modeling on single-cell ATAC-seq data. Nat Methods. 2019 May;16(5):397–400.

Buenrostro JD, Wu B, Litzenburger UM, Ruff D, Gonzales ML, Snyder MP, Chang HY, Greenleaf WJ. Single-cell chromatin accessibility reveals principles of regulatory variation. Nature. 2015 Jul 23;523(7561):486–90.

Carroll SB. Chance and necessity: the evolution of morphological complexity and diversity. Nature 2001;409:1102–9.

Chen X, Miragaia RJ, Natarajan KN, Teichmann SA. A rapid and robust method for single cell chromatin accessibility profiling. Nat Commun. 2018 Dec 17;9(1):5345.

Chin MH, Pellegrini M, Plath K, Lowry WE. Molecular analyses of human induced pluripotent stem cells and embryonic stem cells. Cell Stem Cell. 2010 Aug 6;7(2):263–9.

Christensen CL, Kwiatkowski N, Abraham BJ, Carretero J, Al-Shahrour F, Zhang T, Chipumuro E, Herter-Sprie GS, Akbay EA, Altabef A, Zhang J, Shimamura T, Capelletti M, Reibel JB, Cavanaugh JD, Gao P, Liu Y, Michaelsen SR, Poulsen HS, Aref AR, Barbie DA, Bradner JE, George RE, Gray NS, Young RA, Wong KK. Targeting transcriptional addictions in small cell lung cancer with a covalent CDK7 inhibitor. Cancer Cell. 2014 Dec 8;26(6):909–922.

Cusanovich DA, Daza R, Adey A, Pliner HA, Christiansen L, Gunderson KL, Steemers FJ, Trapnell C, Shendure J. Multiplex single cell profiling of chromatin accessibility by combinatorial cellular indexing. Science. 2015 May 22;348(6237):910–4.

Cusanovich DA, Reddington JP, Garfield DA, Daza RM, Aghamirzaie D, Marco-Ferreres R, Pliner HA, Christiansen L, Qiu X, Steemers FJ, Trapnell C, Shendure J, Furlong EEM. The cis-regulatory dynamics of embryonic development at singlecell resolution. Nature. 2018a Mar 22;555(7697):538–542.

Cusanovich DA, Hill AJ, Aghamirzaie D, Daza RM, Pliner HA, Berletch JB, Filippova GN, Huang X, Christiansen L, DeWitt WS, Lee C, Regalado SG, Read DF, Steemers FJ, Disteche CM, Trapnell C, Shendure J. A Single-Cell Atlas of In Vivo Mammalian Chromatin Accessibility. Cell. 2018b Aug 23;174(5):1309–1324.e18.

Denny SK, Yang D, Chuang CH, Brady JJ, Lim JS, Grüner BM, Chiou SH, Schep AN, Baral J, Hamard C, Antoine M, Wislez M, Kong CS, Connolly AJ, Park KS, Sage J, Greenleaf WJ, Winslow MM. Nfib Promotes Metastasis through a Widespread Increase in Chromatin Accessibility. Cell. 2016 Jul 14;166(2):328–342.

Duan D. Systemic delivery of adeno-associated viral vectors. Curr Opin Virol. 2016 Dec;21:16–25.

ENCODE Project Consortium. An integrated encyclopedia of DNA elements in the human genome. Nature. 2012 Sep 6;489(7414):57–74.

FANTOM Consortium and the RIKEN PMI and CLST (DGT) et al. A promoter-level mammalian expression atlas. Nature. 2014 Mar 27;507(7493):462–70.

Farago AF, Awatramani RB, Dymecki SM. Assembly of the brainstem cochlear nuclear complex is revealed by intersectional and subtractive genetic fate maps. Neuron. 2006 Apr 20;50(2):205–18.

Gao T, He B, Liu S, Zhu H, Tan K, Qian J. EnhancerAtlas: a resource for enhancer annotation and analysis in 105 human cell/tissue types. Bioinformatics. 2016 Dec 1;32(23):3543–3551.

George J, Lim JS, Jang SJ, Cun Y, Ozretić L, Kong G, Leenders F, Lu X, Fernández-Cuesta L, Bosco G, Müller C, Dahmen I, Jahchan NS, Park KS, Yang D, Karnezis AN, Vaka D, Torres A, Wang MS, Korbel JO, Menon R, Chun SM, Kim D, Wilkerson M, Hayes N, Engelmann D, Pützer B, Bos M, Michels S, Vlasic I, Seidel D, Pinther B, Schaub P, Becker C, Altmüller J, Yokota J, Kohno T, Iwakawa R, Tsuta K, Noguchi M, Muley T, Hoffmann H, Schnabel PA, Petersen I, Chen Y, Soltermann A, Tischler V, Choi CM, Kim YH, Massion PP, Zou Y, Jovanovic D, Kontic M, Wright GM, Russell PA, Solomon B, Koch I, Lindner M, Muscarella LA, la-Torre A, Field JK, Jakopovic M, Knezevic J, Castaños-Vélez E, Roz L, Pastorino U, Brustugun OT, Lund-Iversen M, Thunnissen E, Köhler J, Schuler M, Botling J, Sandelin M, Sanchez-Cespedes M, Salvesen HB, Achter V, Lang U, Bogus M, Schneider PM, Zander T, Ansén S, Hallek M, Wolf J, Vingron M, Yatabe Y, Travis WD, Nürnberg P, Reinhardt C, Perner S, Heukamp L, Büttner R, Haas SA, Brambilla E, Peifer M, Sage J, Thomas RK. Comprehensive genomic profiles of small cell lung cancer. Nature. 2015 Aug 6;524(7563):47–53.

Hardee CL, Arévalo-Soliz LM, Hornstein BD, Zechiedrich L. Advances in Non-Viral DNA Vectors for Gene Therapy. Genes (Basel). 2017 Feb 10;8(2). pii: E65.

Hait TA, Amar D, Shamir R, Elkon R. FOCS: a novel method for analyzing enhancer and gene activity patterns infers an extensive enhancer-promoter map. Genome Biol. 2018 May 1;19(1):56.

Hon CC, Shin JW, Carninci P, Stubbington MJT. The Human Cell Atlas: Technical approaches and challenges. Brief Funct Genomics. 2018 Jul 1;17(4):283–294.

Ienasescu H, Li K, Andersson R, Vitezic M, Rennie S, Chen Y, Vitting-Seerup K, Lagoni E, Boyd M, Bornholdt J, de-Hoon MJ, Kawaji H, Lassmann T; FANTOM Consortium, Hayashizaki Y, Forrest AR, Carninci P, Sandelin A. On-the-fly selection of cell-specific enhancers, genes, miRNAs and proteins across the human body using SlideBase. Database (Oxford). 2016 Dec 26;2016. pii: baw144.

Inoue F, Kircher M, Martin B, Cooper GM, Witten DM, McManus MT, Ahituv N, Shendure J. A systematic comparison reveals substantial differences in chromosomal versus episomal encoding of enhancer activity. Genome Res. 2017 Jan;27(1):38–52.

Kron KJ, Bailey SD, Lupien M. Enhancer alterations in cancer: a source for a cell identity crisis. Genome Med. 2014 Sep 23;6(9):77.

Kouno T, Moody J, Kwon AT, Shibayama Y, Kato S, Huang Y, Böttcher M, Motakis E, Mendez M, Severin J, Luginbühl J, Abugessaisa I, Hasegawa A, Takizawa S, Arakawa T, Furuno M, Ramalingam N, West J, Suzuki H, Kasukawa T, Lassmann T, Hon CC, Arner E, Carninci P, Plessy C, Shin JW. C1 CAGE detects transcription start sites and enhancer activity at singlecell resolution. Nat Commun. 2019 Jan 21;10(1):360.

Lakso M, Sauer B, Mosinger-B Jr, Lee EJ, Manning RW, Yu SH, Mulder KL, Westphal H. Targeted oncogene activation by site-specific recombination in transgenic mice. Proc Natl Acad Sci U-S A. 1992 Jul 15;89(14):6232–6.

Lareau C, Kangeyan D, Aryee MJ. Preprocessing and Computational Analysis of Single-Cell Epigenomic Datasets. Methods Mol Biol. 2019;1935:187–202.

Liu Y, Zeng Y, Liu L, Zhuang C, Fu X, Huang W, Cai Z. Synthesizing AND gate genetic circuits based on CRISPR-Cas9 for identification of bladder cancer cells. Nat Commun. 2014 Nov 6;5:5393.

Liu Y, Yu S, Dhiman VK, Brunetti T, Eckart H, White KP. Functional assessment of human enhancer activities using whole-genome STARR-sequencing. Genome Biol. 2017 Nov 20; 18(1):219.

Lizio M, Harshbarger J, Shimoji H, Severin J, Kasukawa T, Sahin S, Abugessaisa I, Fukuda S, Hori F, Ishikawa-Kato S, Mungall CJ, Arner E, Baillie JK, Bertin N, Bono H, de-Hoon M, Diehl AD, Dimont E, Freeman TC, Fujieda K, Hide W, Kaliyaperumal R, Katayama T, Lassmann T, Meehan TF, Nishikata K, Ono H, Rehli M, Sandelin A, Schultes EA, ‘t Hoen PA, Tatum Z, Thompson M, Toyoda T, Wright DW, Daub CO, Itoh M, Carninci P, Hayashizaki Y, Forrest AR, Kawaji H; FANTOM consortium. Gateways to the FANTOM5 promoter level mammalian expression atlas. Genome Biol. 2015 Jan 5;16:22.

Luan H, Peabody NC, Vinson CR, White BH. Refined spatial manipulation of neuronal function by combinatorial restriction of transgene expression. Neuron. 2006 Nov 9;52(3):425–36.

Lukashev AN, Zamyatnin-AA Jr. Viral Vectors for Gene Therapy: Current State and Clinical Perspectives. Biochemistry (Mosc). 2016 Jul;81(7):700–8.

Macosko EZ, Basu A, Satija R, Nemesh J, Shekhar K, Goldman M, Tirosh I, Bialas AR, Kamitaki N, Martersteck EM, Trombetta JJ, Weitz DA, Sanes JR, Shalek AK, Regev A, McCarroll SA. Highly Parallel Genome-wide Expression Profiling of Individual Cells Using Nanoliter Droplets. Cell. 2015 May 21;161(5):1202–14. doi: 10.1016/j.cell.2015.05.002.

Mallo M. Controlled gene activation and inactivation in the mouse. Front Biosci. 2006 Jan 1;11:313–27.

Marei HE, Althani A, Lashen S, Cenciarelli C, Hasan A. Genetically unmatched human iPSC and ESC exhibit equivalent gene expression and neuronal differentiation potential. Sci Rep. 2017 Dec 13;7(1):17504.

Mezger A, Klemm S, Mann I, Brower K, Mir A, Bostick M, Farmer A, Fordyce P, Linnarsson S, Greenleaf W. High-throughput chromatin accessibility profiling at single-cell resolution. Nat Commun. 2018 Sep 7;9(1):3647.

Mora A, Sandve GK, Gabrielsen OS, Eskeland R. In the loop: promoter-enhancer interactions and bioinformatics. Brief Bioinform. 2016 Nov;17(6):980–995.

Morel M, Shtrahman R, Rotter V, Nissim L, Bar-Ziv RH. Cellular heterogeneity mediates inherent sensitivity-specificity tradeoff in cancer targeting by synthetic circuits. Proc Natl Acad Sci U S A. 2016 Jul 19;113(29):8133–8.

Mortazavi A, Pepke S, Jansen C, Marinov GK, Ernst J, Kellis M, Hardison RC, Myers RM, Wold BJ. Integrating and mining the chromatin landscape of cell-type specificity using self-organizing maps. Genome Res. 2013 Dec;23(12):2136–48.

Nissim L, Beatus T, Bar-Ziv R. An autonomous system for identifying and governing a cell’s state in yeast. Phys Biol. 2007 Aug 16;4(3):154–63.

Nissim L, Bar-Ziv RH. A tunable dual-promoter integrator for targeting of cancer cells. Mol Syst Biol. 2010 Dec 21;6:444.

Park KS, Liang MC, Raiser DM, Zamponi R, Roach RR, Curtis SJ, Walton Z, Schaffer BE, Roake CM, Zmoos AF, Kriegel C, Wong KK, Sage J, Kim CF. Characterization of the cell of origin for small cell lung cancer. Cell Cycle. 2011 Aug 15;10(16):2806–15. Epub 2011 Aug 15.

R Core Team (2013). R: A language and environment for statistical computing. R Foundation for Statistical Computing, Vienna, Austria. URL http://www.R-project.org/.

Siuti P, Yazbek J, Lu TK. Synthetic circuits integrating logic and memory in living cells. Nat Biotechnol. 2013 May;31(5):448–52.

Stockinger P, Kvitsiani D, Rotkopf S, Tirián L, Dickson BJ. Neural circuitry that governs Drosophila male courtship behavior. Cell. 2005 Jun 3;121(5):795–807.

Stuart T, Butler A, Hoffman P, Hafemeister C, Papalexi E, Mauck WM 3rd, Hao Y, Stoeckius M, Smibert P, Satija R. Comprehensive Integration of Single-Cell Data. Cell. 2019 Jun 13;177(7):1888–1902.e21.

Suster ML, Seugnet L, Bate M, Sokolowski MB. Refining GAL4-driven transgene expression in Drosophila with a GAL80 enhancer-trap. Genesis. 2004 Aug;39(4):240–5.

Valentine JW, Collins AG, Meyer CP. Morphological complexity increase in metazoans. Paleobiology 1994;20:131–142.

Regev A, Teichmann SA, Lander ES, Amit I, Benoist C, Birney E, Bodenmiller B, Campbell P, Carninci P, Clatworthy M, Clevers H, Deplancke B, Dunham I, Eberwine J, Eils R, Enard W, Farmer A, Fugger L, Göttgens B, Hacohen N, Haniffa M, Hemberg M, Kim S, Klenerman P, Kriegstein A, Lein E, Linnarsson S, Lundberg E, Lundeberg J, Majumder P, Marioni JC, Merad M, Mhlanga M, Nawijn M, Netea M, Nolan G, Pe’er D, Phillipakis A, Ponting CP, Quake S, Reik W, Rozenblatt-Rosen O, Sanes J, Satija R, Schumacher TN, Shalek A, Shapiro E, Sharma P, Shin JW, Stegle O, Stratton M, Stubbington MJT, Theis FJ, Uhlen M, van-Oudenaarden A, Wagner A, Watt F, Weissman J, Wold B, Xavier R, Yosef N; Human Cell Atlas Meeting Participants. The Human Cell Atlas. Elife. 2017 Dec 5;6. pii: e27041.

Struhl G, Basler K. Organizing activity of wingless protein in Drosophila. Cell. 1993 Feb 26;72(4):527–40.

Urrutia E, Chen L, Zhou H, Jiang Y. Destin: toolkit for single-cell analysis of chromatin accessibility. Bioinformatics. 2019 Oct 1;35(19):3818–3820.

Wang X, He L, Goggin SM, Saadat A, Wang L, Sinnott-Armstrong N, Claussnitzer M, Kellis M. High-resolution genome-wide functional dissection of transcriptional regulatory regions and nucleotides in human. Nat Commun. 2018 Dec 19;9(1):5380.

Wickham H. ggplot2: Elegant Graphics for Data Analysis. Springer-Verlag New York, 2016.

Wong JKL, Mohseni R, Hamidieh AA, MacLaren RE, Habib N, Seifalian AM. Limitations in Clinical Translation of Nanoparticle-Based Gene Therapy. Trends Biotechnol. 2017 Dec;35(12):1124–1125.

Xiong L, Xu K, Tian K, Shao Y, Tang L, Gao G, Zhang M, Jiang T, Zhang QC. SCALE method for single-cell ATAC-seq analysis via latent feature extraction. Nat Commun. 2019 Oct 8;10(1):4576.

Zhou W, Ji Z, Fang W, Ji H. Global prediction of chromatin accessibility using small-cell-number and single-cell RNA-seq. Nucleic Acids Res. 2019 Nov 4;47(19):e121.

