## Supplementary Materials for "The intersectional genetics landscape for human"

This file contains Supplementary Figures S1-6 and Supplementary Tables S1-S6, and Supplementary Data S1-2 are available at https://github.com/AndreMacedo88/VEnCode.

**Supplementary Figures**

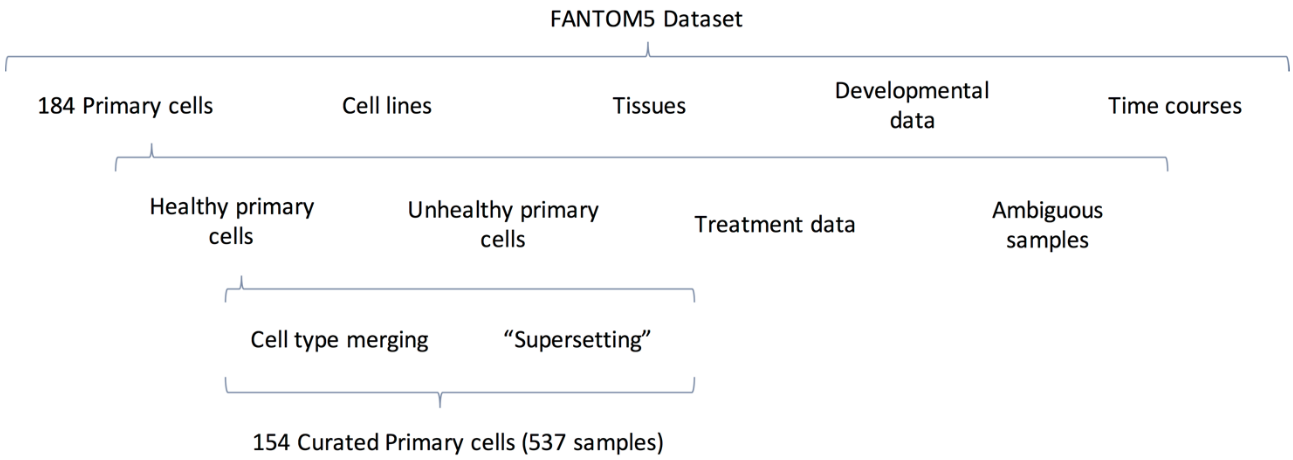


**Figure S1. Pipeline for FANTOM5 data preparation and curation.** Further details on cell type merging, "supersetting", and excluded primary cells are provide Supplementary Table S1.

1. Calculate $E_{raw}$ from various samples of inactive promoters

Vary number cell types →

RE activity

| | |
|---|---|
| *RE1* | 0 0 0 0 0 0 0 0 0 0 0 … |
| *RE2* | 0 0 0 0 0 0 0 0 0 0 0 … |
| … | 0 0 0 0 0 0 0 0 0 0 0 … |
| | 0 0 0 0 0 0 0 0 0 0 0 … |
| | 0 0 0 0 0 0 0 0 0 0 0 … |

Vary $k$

2. Calculate correlation best-fitting curve

Vary number cell types: $y = sk^h$

Coefficient variation with k:

$s = mk + e$   $R^2 = 0.9993$

$h = d + \frac{a - d}{1 + \left(\frac{k}{c}\right)^b}$   $R^2 = 1$

| Coefficients | | | | | |
|---|---|---|---|---|---|
| a | b | c | d | m | e |
| -164054,1 | 0,9998811 | 6,08895E-06 | 1,00051 | 0,9527 | -0,1131 |

3. Generate function that generates expected best $E$

$$y = (mk + e)x^{\left(d + \frac{a-d}{1+\left(\frac{k}{c}\right)^b}\right)}$$

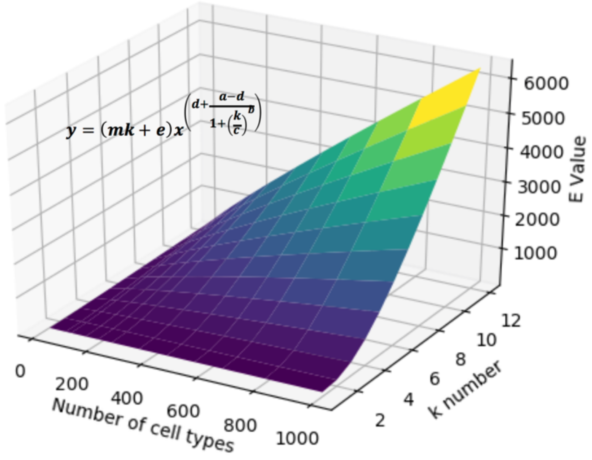


**Figure S2. Generating the function that returns the best possible *E* ($E_{best}$).** We generated $E_{raw}$ (as described in Figure 6) for simulated data reflecting a best-case-scenario for a VEnCode – where all *k* REs are inactive in the non-target cell – varying the number of cell types and *k* in the dataset (1.). By changing cell type number ranging from 20 to 1000, we calculated the best fitting curves that predicted $E_{raw}$ and generated the general equation $y = sk^h$ (2.). However, *s* and *h* depend on the number of REs (*k*). So, varying *k* from 1 up to 10, we obtained the best fitting equations that explain this variation (2.). With this data we then generated a function that best accounts for the variation of both *k* and cell type number in the dataset (3.). The effectiveness of this equation can be seen in Table S2.

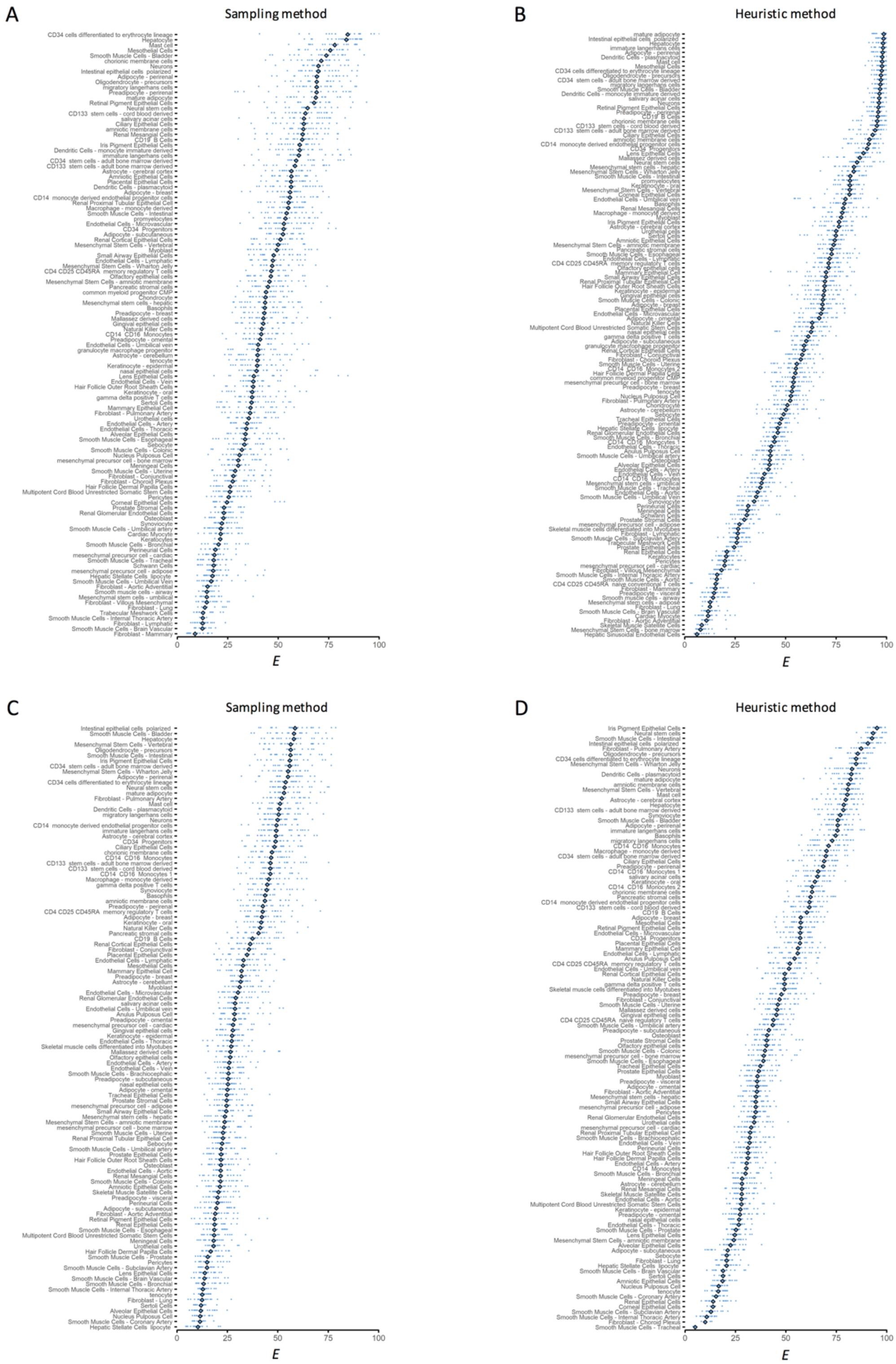


**Figure S3. *E* value variation by cell type.** 5 to 20 VEnCodes at $k$ = 4 were obtained for every possible primary cell type and their $E$ values were determined as described in Figure 6. **A, B.** Results for VEnCodes generated using the promoter dataset. In (**A**) the sampling method (see Figure 3) was used to obtain VEnCodes and determine $E$ for 114 cell types. In **(B)**, the heuristic method (see Figure 4) was used for the same purpose, allowing us to analyze $E$ for up to 20 VEnCodes for 131 cell types. **C, D.** Results for VEnCodes generated using the enhancer dataset. **C.** $E$ value distribution of VEnCodes obtained with the sampling method for 112 primary cell types. **D.** Similarly, 117 cell types had 5-20 enhancer VEnCodes retrieved using the heuristic method and their $E$ was determined.

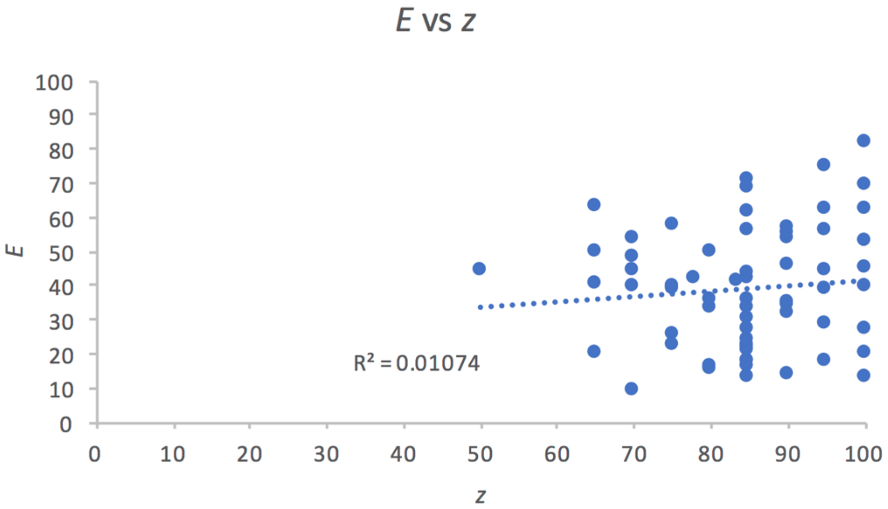


**Figure S4. *E* versus *z* scores.** Plotted is the *E* and *z* scores for each cell type of a list of 64 cell types with three donors and which we managed to retrieve both *E* and *z* values. Also plotted is the linear regression attempt to model the relationship between these two values.

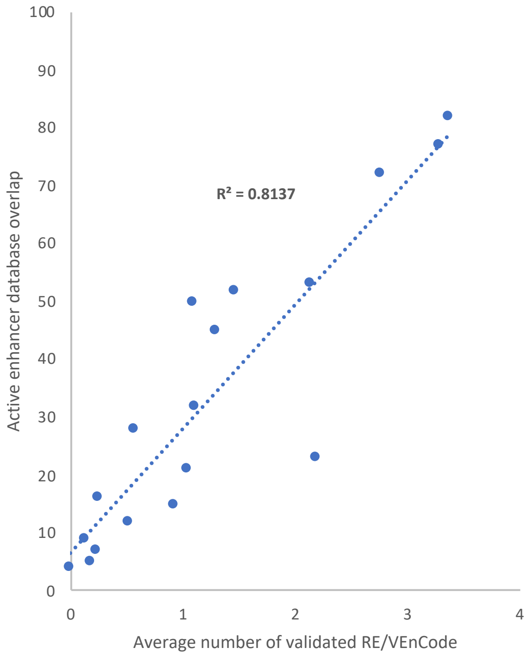


**Figure S5. Correlation between the overlap between external databases and curated CAGE-Seq enhancer databases and the likelihood of VEnCode validation.** The overlap between active enhancers in the curated FANTOM5 CAGE-seq data set and the external data sets used to validate VEnCodes are plotted versus the number of enhancers in a VEnCode of *k* = 4 that are likely to be validated. Each data point represents one of the 18 cell types used in the validation study (Figure 9 and Table S4). 200 VEnCodes were generated from the FANTOM5 data set and validation of every enhancer that composes each VEnCode was accessed as explained in Figure 9A.

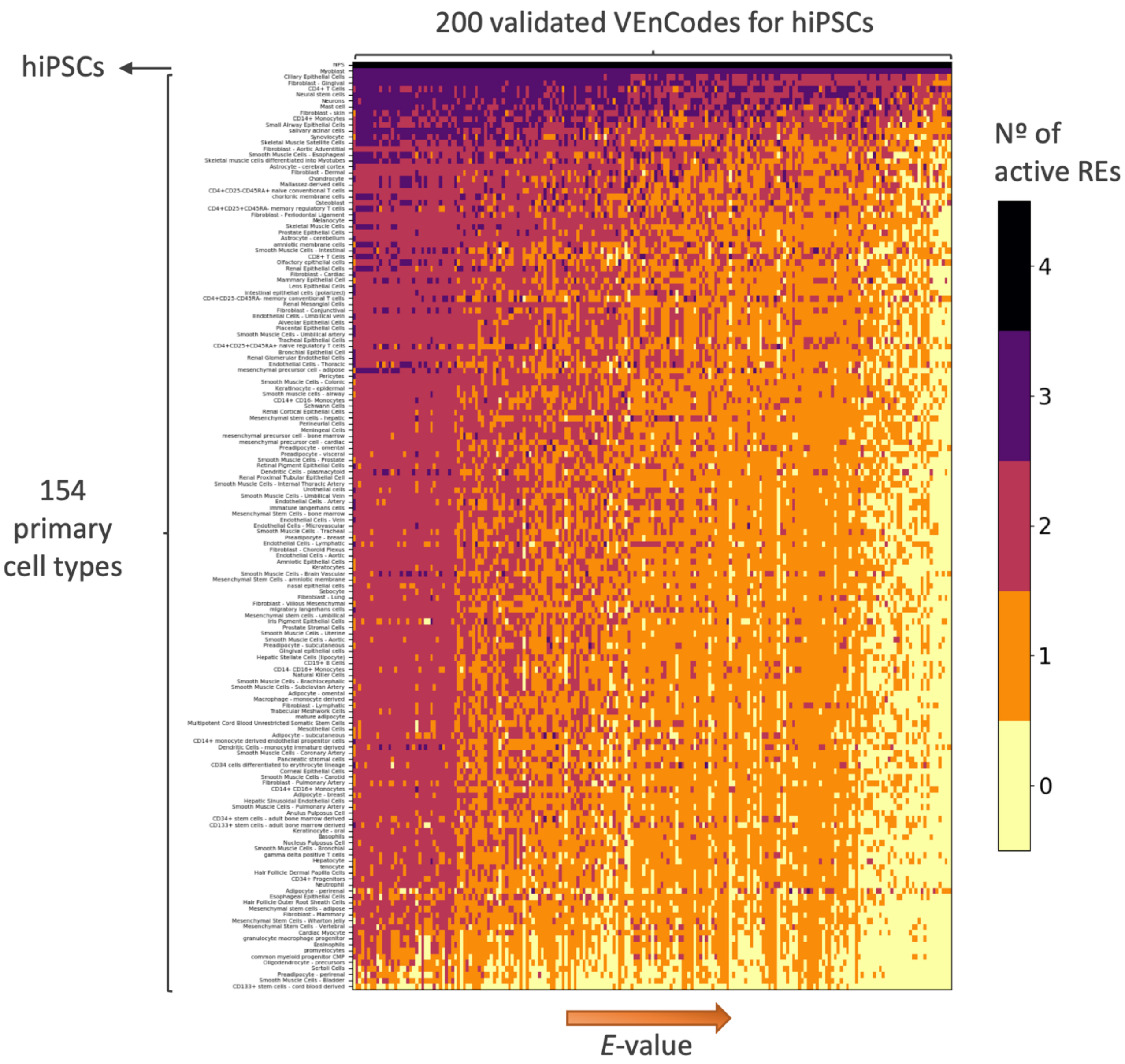


**Figure S6. Validated CAGE-Seq-determined VEnCodes.** Visualization of 200 validated hiPSCs $k$ = 4 VEnCodes generated from enhancers present in both CAGE-seq and ChIP-STARR-seq data sets using the sampling method. Each pixel on the heatmap represents the number of enhancers for each VEnCode (columns), which are active for each cell type (rows). Columns are ordered according to their increasing *E*-value.

### Supplementary Tables

**Table S1. List of curated primary cell types and merged, supersets and excluded categories used in this study.** The list of curated cell types contains 154 primary cell types, encompassing a total of 537 cell line samples. Merging was done with the rationale of turning a vast data on diverse cell conditions into biologically relevant cell types. Merged cell types are then used as curated cell types and the original “cell type” data is not accessed independently. On the other hand, a superset cell type means that the superset data includes the subset data, but each subset is still included in the curated cell type list used in this study. The excluded cell type category lists the data not used at any point throughout the study.

| Curated cell types (154 Primary cell types) | |
|---|---|
| Adipocyte - breast | Mast cell |
| Adipocyte - omental | mature adipocyte |
| Adipocyte - perirenal | Melanocyte |
| Adipocyte - subcutaneous | Meningeal Cells |
| Alveolar Epithelial Cells | mesenchymal precursor cell - adipose |
| Amniotic Epithelial Cells | mesenchymal precursor cell - bone marrow |
| amniotic membrane cells | mesenchymal precursor cell - cardiac |
| Anulus Pulposus Cell | Mesenchymal stem cells - adipose |
| Astrocyte - cerebellum | Mesenchymal Stem Cells - amniotic membrane |
| Astrocyte - cerebral cortex | Mesenchymal Stem Cells - bone marrow |
| Basophils | Mesenchymal stem cells - hepatic |
| Bronchial Epithelial Cell | Mesenchymal stem cells - umbilical |
| Cardiac Myocyte | Mesenchymal Stem Cells - Vertebral |
| CD133+ stem cells - adult bone marrow derived | Mesenchymal Stem Cells - Wharton Jelly |
| CD133+ stem cells - cord blood derived | Mesothelial Cells |
| CD14+ monocyte derived endothelial progenitor cells | migratory langerhans cells |
| CD14+ Monocytes | Multipotent Cord Blood Unrestricted Somatic Stem Cells |
| CD14+CD16- Monocytes | Myoblast |
| CD14+CD16+ Monocytes | nasal epithelial cells |
| CD14-CD16+ Monocytes | Natural Killer Cells |
| CD19+ B Cells | Neural stem cells |
| CD34 cells differentiated to erythrocyte lineage | Neurons |
| CD34+ Progenitors | Neutrophil |
| CD34+ stem cells - adult bone marrow derived | Nucleus Pulposus Cell |
| CD4+ T Cells | Olfactory epithelial cells |
| CD4+CD25+CD45RA- memory regulatory T cells | Oligodendrocyte - precursors |
| CD4+CD25+CD45RA+ naive regulatory T cells | Osteoblast |
| CD4+CD25-CD45RA- memory conventional T cells | Pancreatic stromal cells |
| CD4+CD25-CD45RA+ naive conventional T cells | Pericytes |
| CD8+ T Cells | Perineurial Cells |
| Chondrocyte | Placental Epithelial Cells |
| chorionic membrane cells | Preadipocyte - breast |
| Ciliary Epithelial Cells | Preadipocyte - omental |
| common myeloid progenitor CMP | Preadipocyte - perirenal |
| Corneal Epithelial Cells | Preadipocyte - subcutaneous |
| Dendritic Cells - monocyte immature derived | Preadipocyte - visceral |
| Dendritic Cells - plasmacytoid | promyelocytes |
| Endothelial Cells - Aortic | Prostate Epithelial Cells |
| Endothelial Cells - Artery | Prostate Stromal Cells |
| Endothelial Cells - Lymphatic | Renal Cortical Epithelial Cells |
| Endothelial Cells - Microvascular | Renal Epithelial Cells |
| Endothelial Cells - Thoracic | Renal Glomerular Endothelial Cells |
| Endothelial Cells - Umbilical vein | Renal Mesangial Cells |
| Endothelial Cells - Vein | Renal Proximal Tubular Epithelial Cell |
| Eosinophils | Retinal Pigment Epithelial Cells |
| Esophageal Epithelial Cells | salivary acinar cells |
| Fibroblast - Aortic Adventitial | Schwann Cells |
| Fibroblast - Cardiac | Sebocyte |
| Fibroblast - Choroid Plexus | Sertoli Cells |
| Fibroblast - Conjunctival | Skeletal Muscle Cells |
| Fibroblast - Dermal | Skeletal muscle cells differentiated into Myotubes - multinucleated |
| Fibroblast - Gingival | Skeletal Muscle Satellite Cells |
| Fibroblast - Lung | Small Airway Epithelial Cells |
| Fibroblast - Lymphatic | Smooth muscle cells - airway |
| Fibroblast - Mammary | Smooth Muscle Cells - Aortic |
| Fibroblast - Periodontal Ligament | Smooth Muscle Cells - Bladder |
| Fibroblast - Pulmonary Artery | Smooth Muscle Cells - Brachiocephalic |
| Fibroblast - skin | Smooth Muscle Cells - Brain Vascular |
| Fibroblast - Villous Mesenchymal | Smooth Muscle Cells - Bronchial |
| gamma delta positive T cells | Smooth Muscle Cells - Carotid |
| Gingival epithelial cells | Smooth Muscle Cells - Colonic |
| granulocyte macrophage progenitor | Smooth Muscle Cells - Coronary Artery |
| Hair Follicle Dermal Papilla Cells | Smooth Muscle Cells - Esophageal |
| Hair Follicle Outer Root Sheath Cells | Smooth Muscle Cells - Internal Thoracic Artery |
| Hepatic Sinusoidal Endothelial Cells | Smooth Muscle Cells - Intestinal |
| Hepatic Stellate Cells (lipocyte) | Smooth Muscle Cells - Prostate |
| Hepatocyte | Smooth Muscle Cells - Pulmonary Artery |
| immature langerhans cells | Smooth Muscle Cells - Subclavian Artery |
| Intestinal epithelial cells (polarized) | Smooth Muscle Cells - Tracheal |
| Iris Pigment Epithelial Cells | Smooth Muscle Cells - Umbilical artery |
| Keratinocyte - epidermal | Smooth Muscle Cells - Umbilical Vein |
| Keratinocyte - oral | Smooth Muscle Cells - Uterine |
| Keratocytes | Synoviocyte |
| Lens Epithelial Cells | tenocyte |
| Macrophage - monocyte derived | Trabecular Meshwork Cells |
| Mallassez-derived cells | Tracheal Epithelial Cells |
| Mammary Epithelial Cell | Urothelial cells |

**Table S1. Continued.**

| Merged data | |
|---|---|
| **Merged cell types** | **Original cell types** |
| **CD14+ Monocytes** | CD14+ monocytes - mock treated |
| | CD14+ monocytes - treated with BCG |
| | CD14+ monocytes - treated with B-glucan |
| | CD14+ monocytes - treated with Candida |
| | CD14+ monocytes - treated with Cryptococcus |
| | CD14+ monocytes - treated with Group A streptococci |
| | CD14+ monocytes - treated with IFN + N-hexane |
| | CD14+ monocytes - treated with lipopolysaccharide |
| | CD14+ monocytes - treated with Salmonella |
| | CD14+ monocytes - treated with Trehalose dimycolate (TDM) |
| | CD14+ Monocytes |
| **CD19+ B Cells** | CD19+ B Cells (pluriselect) |
| | CD19+ B Cells |
| **CD4+CD25+CD45RA- memory regulatory T cells** | CD4+CD25+CD45RA- memory regulatory T cells expanded |
| | CD4+CD25+CD45RA- memory regulatory T cells |
| **CD4+CD25+CD45RA+ naive regulatory T cells** | CD4+CD25+CD45RA+ naive regulatory T cells expanded |
| | CD4+CD25+CD45RA+ naive regulatory T cells |
| **CD4+CD25-CD45RA- memory conventional T cells** | CD4+CD25-CD45RA- memory conventional T cells expanded |
| | CD4+CD25-CD45RA- memory conventional T cells |
| **CD4+CD25-CD45RA+ naive conventional T cells** | CD4+CD25-CD45RA+ naive conventional T cells expanded |
| | CD4+CD25-CD45RA+ naive conventional T cells |
| **CD8+ T Cells** | CD8+ T Cells (pluriselect) |
| | CD8+ T Cells |
| **Chondrocyte** | Chondrocyte - de diff |
| | Chondrocyte - re diff |
| **Fibroblast - skin** | Fibroblast - skin dystrophia myotonica |
| | Fibroblast - skin normal |
| | Fibroblast - skin spinal muscular atrophy |
| | Fibroblast - skin walker warburg |
| **Mast cell** | Mast cell - stimulated |
| | Mast cell |
| **Melanocyte** | Melanocyte - dark |
| | Melanocyte - light |
| **Neutrophil** | neutrophil PMN |
| | Neutrophils |
| **Prostate Epithelial Cells** | Prostate Epithelial Cells (polarized) |
| | Prostate Epithelial Cells |

**Table S1. Continued.**

| Curated supersets: superset | Curated supersets: subset | Excluded cell types |
|---|---|---|
| **CD14+ Monocytes** | CD14+CD16- Monocytes | mesenchymal precursor cell - ovarian cancer left ovary |
| | CD14+CD16+ Monocytes | mesenchymal precursor cell - ovarian cancer metastasis |
| | | mesenchymal precursor cell - ovarian cancer right ovary |
| **CD4+ T Cells** | CD4+CD25+CD45RA- memory regulatory T cells | Osteoblast - differentiated |
| | CD4+CD25+CD45RA+ naive regulatory T cells | Peripheral Blood Mononuclear Cells |
| | CD4+CD25-CD45RA- memory conventional T cells | Whole blood (ribopure) |
| | CD4+CD25-CD45RA+ naive conventional T cells | |

**Table S2. Normalized *E* (*E* = $E_{raw}/E_{best}$) values for a range of number of cell types in data and *k* REs used to generate a VEnCode.** Values were generated, as described in Figure 6, for a intraindividual robust best-case possible VEnCode and normalized using the function described in Figure S2. Thus, the values reflect the average *E* expected for the most intraindividual robust VEnCodes.

| | | Number of Cell types | | | | | | | | | | | |
|---|---|---|---|---|---|---|---|---|---|---|---|---|---|
| | | **20** | **80** | **100** | **154** | **200** | **250** | **350** | **450** | **550** | **650** | **800** | **1000** |
| *k* | **1** | 100.0 | 100.0 | 100.0 | 100.0 | 100.0 | 100.0 | 100.0 | 100.0 | 100.0 | 100.0 | 100.0 | 100.0 |
| | **2** | 99.5 | 98.8 | 96.5 | 96.1 | 97.9 | 100.0 | 97.2 | 97.3 | 97.4 | 100.0 | 100.0 | 100.0 |
| | **3** | 96.3 | 98.9 | 97.7 | 96.7 | 98.6 | 97.9 | 95.8 | 98.6 | 95.2 | 97.1 | 99.5 | 95.9 |
| | **4** | 99.2 | 98.4 | 97.3 | 98.4 | 97.6 | 98.4 | 99.1 | 98.3 | 98.5 | 98.6 | 98.7 | 97.0 |
| | **5** | 99.5 | 98.9 | 99.1 | 99.1 | 97.7 | 99.0 | 99.0 | 97.2 | 98.8 | 98.5 | 99.1 | 99.0 |
| | **6** | 98.5 | 99.9 | 99.4 | 99.3 | 98.5 | 99.7 | 99.3 | 99.4 | 99.1 | 99.0 | 99.1 | 99.8 |
| | **7** | 100.0 | 99.9 | 100.0 | 98.9 | 99.5 | 99.9 | 100.0 | 99.3 | 99.7 | 99.2 | 98.0 | 100.0 |
| | **8** | 99.9 | 100.0 | 100.0 | 100.0 | 100.0 | 100.0 | 100.0 | 100.0 | 100.0 | 100.0 | 100.0 | 99.6 |
| | **9** | 100.0 | 100.0 | 100.0 | 99.9 | 100.0 | 100.0 | 100.0 | 99.9 | 100.0 | 100.0 | 100.0 | 100.0 |
| | **10** | 100.0 | 100.0 | 100.0 | 99.7 | 100.0 | 100.0 | 100.0 | 100.0 | 100.0 | 100.0 | 100.0 | 100.0 |

**Table S3. List of curated cancer cell types used in this study.** The list contains 158 biologically relevant cell types, encompassing a total of 274 cancer cell line samples.

| **Curated cancer cell types** | |
|---|---|
| acantholytic squamous carcinoma cell line:HCC1806 | large cell non-keratinizing squamous carcinoma cell line:SKG-II-SF |
| acute lymphoblastic leukemia (B-ALL) cell line | leiomyoblastoma cell line:G-402 |
| acute lymphoblastic leukemia (T-ALL) cell line | leiomyoma cell line |
| acute myeloid leukemia (FAB M0) cell line | leiomyosarcoma cell line:Hs 5 |
| acute myeloid leukemia (FAB M1) cell line | lens epithelial cell line:SRA |
| acute myeloid leukemia (FAB M2) cell line | liposarcoma cell line |
| acute myeloid leukemia (FAB M3) cell line | lung adenocarcinoma cell line |
| acute myeloid leukemia (FAB M4) cell line | lung adenocarcinoma papillary cell line:NCI-H441 |
| acute myeloid leukemia (FAB M4eo) cell line | lymphangiectasia cell line:DS-1 |
| acute myeloid leukemia (FAB M5) cell line | lymphoma malignant hairy B-cell cell line:MLMA |
| acute myeloid leukemia (FAB M6) cell line | malignant trichilemmal cyst cell line:DJM-1 |
| acute myeloid leukemia (FAB M7) cell line | maxillary sinus tumor cell line:HSQ-89 |
| adenocarcinoma cell line:IM95m | medulloblastoma cell line |
| adrenal cortex adenocarcinoma cell line:SW-13 | melanoma cell line |
| adult T-cell leukemia cell line:ATN-1 | meningioma cell line:HKBMM |
| alveolar cell carcinoma cell line:SW 1573 | merkel cell carcinoma cell line |
| anaplastic carcinoma cell line:8305C | mesenchymal stem cell line:Hu5/E18 |
| anaplastic large cell lymphoma cell line:Ki-JK | mesodermal tumor cell line:HIRS-BM |
| anaplastic squamous cell carcinoma cell line:RPMI 2650 | Epithelioid mesothelioma cell line |
| argyrophil small cell carcinoma cell line:TC-YIK | Sarcomatoid mesothelioma cell line |
| astrocytoma cell line:TM-31 | Biphasic mesothelioma cell line |
| b cell line:RPMI1788 | mixed mullerian tumor cell line:HTMMT |
| B lymphoblastoid cell line: GM12878 ENCODE | mucinous adenocarcinoma cell line:JHOM-1 |
| basal cell carcinoma cell line:TE 354.T | mucinous cystadenocarcinoma cell line:MCAS |
| bile duct carcinoma cell line | myelodysplastic syndrome cell line:SKM-1 |
| biphenotypic B myelomonocytic leukemia cell line:MV-4-11 | myeloma cell line:PCM6 |
| bone marrow stromal cell line:StromaNKtert | myxofibrosarcoma cell line |
| breast carcinoma cell line | neuroblastoma cell line |
| bronchial squamous cell carcinoma cell line:KNS-62 | neuroectodermal tumor cell line |
| bronchioalveolar carcinoma cell line | neuroepithelioma cell line:SK-N-MC |
| bronchogenic carcinoma cell line:ChaGo-K-1 | neurofibroma cell line:Hs 53 |
| Burkitt lymphoma cell line | NK T cell leukemia cell line:KHYG-1 |
| carcinoid cell line | non T non B acute lymphoblastic leukemia cell line:P30/OHK |
| carcinosarcoma cell line:JHUCS-1 | non-small cell lung cancer cell line:NCI-H1385 |
| cervical cancer cell line | normal embryonic palatal mesenchymal cell line:HEPM |
| cholangiocellular carcinoma cell line:HuH-28 | normal intestinal epithelial cell line:FHs 74 Int |
| chondrosarcoma cell line:SW 1353 | oral squamous cell carcinoma cell line |
| choriocarcinoma cell line | osteoclastoma cell line:Hs 706 |
| chronic lymphocytic leukemia cell line:SKW-3 | osteosarcoma cell line |
| chronic megakaryoblastic cell line:MEG-01 | pagetoid sarcoma cell line:Hs 925 |
| chronic myeloblastic leukemia cell line:KCL-22 | pancreatic carcinoma cell line:NOR-P1 |
| chronic myelogenous leukemia cell line | papillary adenocarcinoma cell line:8505C |
| clear cell carcinoma cell line | papillotubular adenocarcinoma cell line:TGBC18TKB |
| colon carcinoma cell line | peripheral neuroectodermal tumor cell line:KU-SN |
| cord blood derived cell line:COBL-a untreated | pharyngeal carcinoma cell line:Detroit 562 |
| diffuse large B-cell lymphoma cell line:CTB-1 | plasma cell leukemia cell line:ARH-77 |
| ductal cell carcinoma cell line | pleomorphic hepatocellular carcinoma cell line:SNU-387 |
| embryonic kidney cell line: HEK293/SLAM untreated | prostate cancer cell line |
| embryonic pancreas cell line | rectal cancer cell line:TT1TKB |
| endometrial carcinoma cell line:OMC-2 | renal cell carcinoma cell line |
| endometrial stromal sarcoma cell line:OMC-9 | retinoblastoma cell line:Y79 |
| endometrioid adenocarcinoma cell line:JHUEM-1 | rhabdomyosarcoma cell line |
| epidermoid carcinoma cell line | sacrococcigeal teratoma cell line:HTST |
| epithelioid sarcoma cell line | schwannoma cell line:HS-PSS |
| epitheloid carcinoma cell line: HelaS3 ENCODE | serous adenocarcinoma cell line |
| Ewing sarcoma cell line:Hs 863 | serous cystadenocarcinoma cell line:HTOA |
| extraskeletal myxoid chondrosarcoma cell line:H-EMC-SS | signet ring carcinoma cell line |
| fibrosarcoma cell line:HT-1080 | small cell cervical cancer cell line:HCSC-1 |
| fibrous histiocytoma cell line:GCT TIB-223 | small cell gastrointestinal carcinoma cell line:ECC10 |
| gall bladder carcinoma cell line | small cell lung carcinoma cell line |
| gastric adenocarcinoma cell line | small-cell gastrointestinal carcinoma cell line:ECC4 |
| gastric cancer cell line | somatostatinoma cell line:QGP-1 |
| gastrointestinal carcinoma cell line:ECC12 | spindle cell sarcoma cell line:Hs 132 |
| giant cell carcinoma cell line | splenic lymphoma with villous lymphocytes cell line:SLVL |
| glassy cell carcinoma cell line:HOKUG | squamous cell carcinoma cell line:EC-GI-10.CNhs11252.10463-106H4 |
| glioblastoma cell line | squamous cell carcinoma cell line:JHUS-nk1.CNhs11749.10646-109A7 |
| glioma cell line:GI-1 | squamous cell carcinoma cell line:T3M-5.CNhs11739.10616-108G4 |
| granulosa cell tumor cell line:KGN | squamous cell lung carcinoma cell line |
| hairy cell leukemia cell line:Mo | synovial sarcoma cell line:HS-SY-II |
| Hep-2 cells mock treated | T cell lymphoma cell line:HuT 102 TIB-162 |
| hepatic mesenchymal tumor cell line:LI90 | teratocarcinoma cell line |
| hepatoblastoma cell line:HuH-6 | testicular germ cell embryonal carcinoma cell line |
| hepatocellular carcinoma cell line: HepG2 ENCODE | thymic carcinoma cell line:Ty-82 |
| hepatoma cell line:Li-7 | thyroid carcinoma cell line |
| hereditary spherocytic anemia cell line:WIL2-NS | transitional cell carcinoma cell line |
| Hodgkin lymphoma cell line:HD-Mar2 | tridermal teratoma cell line:HGRT |
| keratoacanthoma cell line:HKA-1 | tubular adenocarcinoma cell line:SUIT-2 |
| Krukenberg tumor cell line:HSKTC | Wilms tumor cell line |
| large cell lung carcinoma cell line | xeroderma pigentosum b cell line:XPL 17 |

**Table S4. Cell types used for the cross-validation studies.** Full references are provided in the main text. Notice that the small cell lung carcinoma cell (SCLC) line H82 was also analyzed separately from the other SCLC lines as there was a separate database (ChIP-seq data from Christensen *et al.,* 2019) that allowed its validation as a single SCLC line, as depicted below.

| Cell type in CAGE | Cell type in external datasets | Type | Methods for RE activity estimation | References |
|---|---|---|---|---|
| Human induced pluripotent stem cell (hiPSC) | human embryonic stem cell (hESC) | pluripotent stem cell | ChIP-STARR-seq | Barakat *et al*., 2018 |
| CD19+ B cells | CD19+ cells | primary cells | TF Binding, eRNA, Histone modifications | Gao *et al*., 2016 |
| CD34+ stem cells - adult bone marrow derived | CD34+ cells | primary cells | TF Binding, DHS, Histone modifications | Gao *et al*., 2016 |
| lung adenocarcinoma cell line: A549 | A549 | cell lines | TF Binding, DHS, FAIRE, eRNA, EP300, POL2, Histone modifications | Gao *et al*., 2016 |
| acute myeloid leukemia (FAB M3) cell line: HL60 | HL60 | cell lines | TF Binding, DHS, eRNA, POL2 | Gao *et al*., 2016 |
| colon carcinoma cell line: CACO-2 | CACO-2 | cell lines | TF Binding, DHS, eRNA | Gao *et al*., 2016 |
| B lymphoblastoid cell line: GM12878 ENCODE | GM12878 | cell lines | ATAC-STARR-seq, TF Binding, DHS, FAIRE, eRNA, EP300, POL2, Histone modifications | Gao *et al*., 2016, Wang *et al*., 2018 |
| small cell lung carcinoma cell line: H82 | H82 | cell lines | ChIP-seq | Christensen *et al*., 2014 |
| hepatocellular carcinoma cell line: HepG2 ENCODE | HepG2 | cell lines | ChIP-seq, TF Binding, DHS, FAIRE, eRNA, EP300, POL2, Histone modifications | Gao *et al*., 2016, Inoue *et al*., 2017 |
| acute lymphoblastic leukemia (T-ALL) cell line: Jurkat | Jurkat | cell lines | TF Binding, DHS, eRNA, Histone modifications | Gao *et al*., 2016 |
| chronic myelogenous leukemia cell line: K562 | K562 | cell lines | TF Binding, DHS, FAIRE, eRNA, EP300, POL2, ChIA-PET, Histone modifications | Gao *et al*., 2016 |
| breast carcinoma cell line: MCF7 | MCF7 | cell lines | TF Binding, DHS, FAIRE, eRNA, EP300, POL2, ChIA-PET, Histone modifications | Gao *et al*., 2016 |
| prostate cancer cell lines: DU145 and PC-3 | prostate cancer cell line: LNCaP | cell lines | WHG-STARR-seq | Liu *et al*., 2017 |
| Burkitt's lymphoma cell line: RAJI | Raji | cell lines | DHS, eRNA, POL2 | Gao *et al*., 2016 |
| small cell lung carcinoma cell lines: DMS144, LK-2, H82,WA-hT | small cell lung carcinoma cell lines: H69, H82, GLC16, H14, H29, H52, H88 | cell lines | ATAC-seq | Christensen *et al*., 2014, Denny *et al*., 2017 |
| neuroepithelioma cell line: SK-N-MC | SK-N-MC | cell lines | TF Binding, DHS, eRNA, POL2 | Gao *et al*., 2016 |
| cervix adenocarcinoma epithelial cell line: HeLa | HeLaS3 | cell lines | TF Binding, DHS, FAIRE, eRNA, EP300, Histone modifications | Gao *et al*., 2016 |
| transformed embryonic kidney cell line: HEK293-SLAM | HEK293 | cell lines | DHS, eRNA, EP300, POL2 | Gao *et al*., 2016 |

**Table S5. List of 35 cells (A549 cell line) retrieved from Kuono *et al*., 2019.**

| Cell Codes |
|---|
| CAGE_4_A04 |
| CAGE_4_A05 |
| CAGE_4_A07 |
| CAGE_4_A09 |
| CAGE_4_B01 |
| CAGE_4_C01 |
| CAGE_4_C06 |
| CAGE_4_C12 |
| CAGE_4_D11 |
| CAGE_4_D12 |
| CAGE_4_E03 |
| CAGE_4_E04 |
| CAGE_4_E05 |
| CAGE_4_E08 |
| CAGE_4_F05 |
| CAGE_4_F07 |
| CAGE_4_F12 |
| CAGE_4_H03 |
| CAGE_4_H07 |
| CAGE_5_A01 |
| CAGE_5_A08 |
| CAGE_5_B10 |
| CAGE_5_C04 |
| CAGE_5_E02 |
| CAGE_5_H10 |
| CAGE_6_D03 |
| CAGE_6_E06 |
| CAGE_6_F08 |
| CAGE_6_F12 |
| CAGE_6_G09 |
| CAGE_6_H07 |
| CAGE_6_H09 |
| CAGE_6_H10 |
| CAGE_6_H11 |
| CAGE_6_H12 |

**Table S6. List of 540 enhancers active in A549 cells as retrieved from the normalized active enhancer calls obtained from C1 CAGE-seq data from the 35 cells described above from Kuono *et al.*, 2019.**

| Enhancers | | | | |
|---|---|---|---|---|
| chr1:100859593-100859940 | chr4:140982751-140983079 | chr8:6323096-6323426 | chr12:47474423-47475249 | chr18:56339458-56339877 |
| chr1:109499781-109500180 | chr4:141011416-141011614 | chr8:72753377-72753463 | chr12:47701468-47701752 | chr18:71010965-71011298 |
| chr1:112196411-112196803 | chr4:141015177-141015539 | chr8:98865970-98866240 | chr12:52433623-52433924 | chr18:74171783-74172228 |
| chr1:117132073-117132239 | chr4:141071687-141072625 | chr9:109932773-109933133 | chr12:52674043-52674518 | chr18:74204839-74205357 |
| chr1:144533627-144534178 | chr4:146546923-146547210 | chr9:109933165-109933377 | chr12:55806634-55807051 | chr19:12888910-12889352 |
| chr1:149816287-149817606 | chr4:151641898-151642513 | chr9:118917854-118918405 | chr12:55807914-55808439 | chr19:13262107-13262928 |
| chr1:15519488-15519688 | chr4:184423820-184424211 | chr9:125168700-125168938 | chr12:55808460-55808880 | chr19:14732511-14733275 |
| chr1:156473111-156473378 | chr4:190942271-190942591 | chr9:132257886-132258547 | chr12:63149457-63149757 | chr19:16188924-16190024 |
| chr1:157030350-157030742 | chr4:24660007-24660225 | chr9:132330057-132330508 | chr12:66628292-66629290 | chr19:18485152-18485556 |
| chr1:159892447-159892988 | chr4:38859134-38859762 | chr9:132369856-132370376 | chr12:69036661-69037118 | chr19:34850085-34850642 |
| chr1:168338134-168338331 | chr4:88128179-88128500 | chr9:134573283-134574205 | chr12:69634325-69634686 | chr19:35491833-35492367 |
| chr1:171652961-171653360 | chr4:88128695-88128755 | chr9:136236880-136237291 | chr12:7592182-7592797 | chr19:35494139-35494542 |
| chr1:192216449-192216576 | chr5:105752390-105752961 | chr9:136739075-136739598 | chr12:94817703-94818348 | chr19:39411140-39411539 |
| chr1:198889384-198889935 | chr5:114726097-114726311 | chr9:140023869-140024391 | chr13:109921804-109922211 | chr19:41681944-41682199 |
| chr1:203481982-203482534 | chr5:114847596-114847935 | chr9:18718417-18718817 | chr13:110790027-110791241 | chr19:41819266-41819494 |
| chr1:206859748-206860259 | chr5:132576270-132576709 | chr9:19161675-19162031 | chr13:111044675-111045176 | chr19:41831643-41832527 |
| chr1:211035320-211035533 | chr5:134259128-134259903 | chr9:21695810-21696232 | chr13:111049993-111050394 | chr19:42756723-42757090 |
| chr1:211988342-211989080 | chr5:134261311-134262635 | chr9:22216803-22217265 | chr13:24133122-24133556 | chr19:45195366-45195649 |
| chr1:220427006-220427444 | chr5:138262166-138262549 | chr9:2875811-2876206 | chr13:24697003-24697328 | chr19:45228436-45228775 |
| chr1:224180261-224180516 | chr5:139031397-139032095 | chr9:35491656-35492052 | chr13:42269057-42269475 | chr19:45242034-45242469 |
| chr1:228782602-228782886 | chr5:139568486-139568801 | chr9:35492185-35492605 | chr13:42646001-42646399 | chr19:45248499-45249479 |
| chr1:234746093-234747674 | chr5:153939934-153940118 | chr9:35971862-35972364 | chr13:49761885-49761908 | chr19:45350775-45351289 |
| chr1:23888678-23889201 | chr5:15451801-15451916 | chr9:37835930-37836292 | chr13:67674730-67675228 | chr19:45352951-45353140 |
| chr1:23903714-23904323 | chr5:154871450-154872048 | chr9:68455261-68455556 | chr14:102414226-102414964 | chr19:45953764-45954783 |
| chr1:240306165-240306375 | chr5:167909535-167909826 | chr9:75518800-75519107 | chr14:103539359-103539722 | chr19:46683956-46684315 |
| chr1:240306412-240306787 | chr5:171681735-171682053 | chr9:90117444-90117744 | chr14:22080552-22080764 | chr19:47363167-47364689 |
| chr1:240306890-240307603 | chr5:17302742-17303327 | chr9:91795136-91795669 | chr14:35628271-35628433 | chr19:47613231-47613789 |
| chr1:240307621-240308162 | chr5:173193377-173193650 | chr9:97684288-97685133 | chr14:50328882-50329315 | chr19:47776748-47776969 |
| chr1:240308207-240308537 | chr5:176359562-176359884 | chr9:97852724-97853044 | chr14:64180685-64181078 | chr19:47886542-47886975 |
| chr1:243431437-243431793 | chr5:25910581-25911176 | chr9:97881296-97881429 | chr14:68555905-68556402 | chr19:48642255-48642637 |
| chr1:24349340-24349488 | chr5:34585438-34585952 | chr10:103124623-103124787 | chr14:69424468-69424783 | chr19:49836983-49837445 |
| chr1:27175784-27176273 | chr5:34586139-34586737 | chr10:103124787-103125187 | chr14:71797093-71797681 | chr19:53727403-53727861 |
| chr1:27527451-27527688 | chr5:58423311-58423752 | chr10:103453403-103453880 | chr14:75086409-75086922 | chr19:55627475-55627790 |
| chr1:33077721-33078012 | chr5:58866470-58866828 | chr10:105519063-105519154 | chr14:77499482-77500543 | chr19:56117286-56117783 |
| chr1:33802277-33802571 | chr5:66178568-66178972 | chr10:112220227-112220521 | chr14:81685513-81686063 | chr19:58912594-58912944 |
| chr1:37944084-37944971 | chr5:82576471-82576688 | chr10:112261043-112261328 | chr14:93540905-93541183 | chr19:6801116-6801467 |
| chr1:43388597-43388971 | chr5:95295681-95296335 | chr10:112263780-112264181 | chr14:99439262-99439638 | chr19:751766-752111 |
| chr1:51245588-51245972 | chr6:11210069-11210603 | chr10:120692616-120692979 | chr15:29526240-29526874 | chr19:782330-783005 |
| chr1:65383044-65383557 | chr6:121759226-121759877 | chr10:123873239-123873409 | chr15:33158575-33159178 | chr19:8674150-8674979 |
| chr1:68257383-68257657 | chr6:127980874-127981174 | chr10:128943463-128943803 | chr15:41953918-41954236 | chr20:1258531-1258969 |
| chr1:77165935-77166101 | chr6:147124877-147125201 | chr10:133879247-133879524 | chr15:52153526-52153874 | chr20:2855066-2855678 |
| chr1:78557346-78557710 | chr6:14777704-14777964 | chr10:13692317-13692641 | chr15:63795379-63795825 | chr20:30301047-30301377 |
| chr1:8933581-8933925 | chr6:150200694-150201094 | chr10:13908290-13908478 | chr15:64540216-64540468 | chr20:32803456-32803920 |
| chr2:109201604-109202307 | chr6:151366985-151367228 | chr10:21317807-21317920 | chr15:65596702-65597188 | chr20:33861234-33861630 |
| chr2:110871797-110872011 | chr6:166364914-166365167 | chr10:32735821-32736068 | chr15:83680959-83681167 | chr20:36772576-36772945 |
| chr2:118740185-118740994 | chr6:166378442-166378825 | chr10:33269452-33269916 | chr15:89532902-89533939 | chr20:3788108-3788452 |
| chr2:127975624-127975823 | chr6:166422302-166422862 | chr10:33272272-33272613 | chr15:90365179-90365539 | chr20:42828501-42828745 |
| chr2:128598749-128599093 | chr6:25992254-25993073 | chr10:3848037-3849536 | chr15:99193577-99194531 | chr20:47750965-47751079 |
| chr2:153277978-153278373 | chr6:29932777-29932958 | chr10:44100382-44100859 | chr16:14395683-14396371 | chr20:48311078-48311758 |
| chr2:157190328-157191531 | chr6:30716597-30717062 | chr10:47057375-47057652 | chr16:14397958-14398068 | chr20:50366772-50367203 |
| chr2:191709136-191709198 | chr6:30737456-30737900 | chr10:47080536-47081217 | chr16:15583408-15583655 | chr20:52239995-52240318 |
| chr2:200209653-200209909 | chr6:31008975-31009347 | chr10:5045021-5045414 | chr16:22082738-22082854 | chr20:52784442-52785020 |
| chr2:201995421-201995893 | chr6:51275260-51275721 | chr10:51488946-51489392 | chr16:24855986-24856230 | chr20:56251701-56252046 |
| chr2:204499475-204499823 | chr6:53376137-53376766 | chr10:516106-516519 | chr16:30645556-30645885 | chr20:56255217-56255759 |
| chr2:222826690-222827411 | chr6:64151294-64151779 | chr10:5333644-5334212 | chr16:31264985-31265278 | chr20:56270075-56270492 |
| chr2:225133197-225133515 | chr6:72092992-72093394 | chr10:5587810-5588190 | chr16:377235-377950 | chr20:56273255-56273697 |
| chr2:225392496-225392764 | chr6:80663174-80663336 | chr10:5783071-5783414 | chr16:68271348-68271876 | chr20:56293544-56293843 |
| chr2:238421741-238422134 | chr6:80777050-80777287 | chr10:74056833-74058104 | chr16:68320188-68320357 | chr20:5637936-5638327 |
| chr2:240196255-240196671 | chr6:94545083-94545465 | chr10:74080430-74080993 | chr16:69790083-69790481 | chr20:56592800-56593208 |
| chr2:241374162-241374480 | chr7:101775527-101775981 | chr10:80925867-80926207 | chr16:69790577-69791500 | chr20:60934143-60934390 |
| chr2:26220678-26220902 | chr7:105922869-105923204 | chr10:92921990-92922253 | chr16:73104505-73104770 | chr20:62494990-62495823 |
| chr2:3340825-3341028 | chr7:107573965-107574445 | chr10:95225325-95225770 | chr16:74701397-74701877 | chr20:641092-641461 |
| chr2:36584242-36585529 | chr7:128696279-128696583 | chr10:98332019-98332237 | chr16:81465654-81465943 | chr21:11174514-11174910 |
| chr2:36790527-36791707 | chr7:130593505-130595916 | chr11:110326620-110326859 | chr16:85734545-85734806 | chr21:18891260-18891546 |
| chr2:37885188-37885619 | chr7:130601178-130601963 | chr11:122888881-122889273 | chr16:87886258-87886670 | chr21:30374865-30375265 |
| chr2:42046531-42047126 | chr7:130920397-130920794 | chr11:125937888-125938141 | chr16:87899593-87899990 | chr21:36208387-36209191 |
| chr2:46931823-46932125 | chr7:134117156-134117311 | chr11:128376891-128377234 | chr17:20688064-20688832 | chr21:36214787-36216208 |
| chr2:58334895-58335527 | chr7:138123858-138124244 | chr11:34331523-34331863 | chr17:20690369-20690534 | chr21:38935728-38936411 |
| chr2:58339709-58340405 | chr7:148684338-148684706 | chr11:3443739-3443990 | chr17:2302778-2303366 | chr21:38936686-38937128 |
| chr2:60962204-60962802 | chr7:152160715-152161193 | chr11:34622320-34622551 | chr17:33787408-33787803 | chr21:44761192-44761567 |
| chr2:61991278-61991660 | chr7:1572262-1572844 | chr11:34676467-34676782 | chr17:35292129-35292558 | chr21:44859375-44859527 |
| chr2:68472906-68473226 | chr7:158224670-158224991 | chr11:46264726-46265198 | chr17:35446954-35447324 | chr21:44914466-44914931 |
| chr2:69989990-69990667 | chr7:22609345-22610034 | chr11:56677602-56677921 | chr17:38601076-38601487 | chr21:44918002-44918401 |
| chr2:739928-740199 | chr7:2875013-2875360 | chr11:61523945-61524112 | chr17:38655848-38656036 | chr21:45020604-45020960 |
| chr2:86922220-86922584 | chr7:42927750-42928726 | chr11:61738760-61739751 | chr17:38689625-38689996 | chr21:45146832-45147391 |
| chr2:91954549-91954973 | chr7:4681329-4681565 | chr11:63949132-63949273 | chr17:38707495-38707811 | chr21:46097395-46097576 |
| chr2:91960340-91960566 | chr7:47510195-47510559 | chr11:65184888-65185395 | chr17:38708057-38709216 | chr22:22011632-22012115 |
| chr2:92317324-92317742 | chr7:47611156-47611689 | chr11:65187297-65188198 | chr17:43305165-43306978 | chr22:25082452-25082825 |
| chr2:92323832-92324227 | chr7:4765280-4765661 | chr11:65254949-65255610 | chr17:43385085-43385291 | chr22:28114157-28114741 |
| chr3:141131755-141132151 | chr7:4778448-4778853 | chr11:65259930-65260851 | chr17:48105016-48105270 | chr22:30601535-30602099 |
| chr3:14444961-14445761 | chr7:5594554-5594977 | chr11:65666492-65667203 | chr17:48228924-48229616 | chr22:30649196-30649657 |
| chr3:148704982-148705163 | chr7:5634980-5635411 | chr11:67037275-67037527 | chr17:48255388-48256117 | chr22:30938421-30938985 |
| chr3:149657313-149657850 | chr7:75567331-75567651 | chr11:67138852-67139479 | chr17:48367346-48367814 | chr22:41460471-41460696 |
| chr3:156432757-156433079 | chr7:84613691-84613734 | chr11:76480789-76480971 | chr17:56736656-56737064 | chr22:41809427-41810696 |
| chr3:156527707-156527883 | chr8:124248679-124249062 | chr11:77445522-77445800 | chr17:57698412-57698732 | chr22:45944996-45945587 |
| chr3:15780032-15780074 | chr8:126231490-126231859 | chr11:90016507-90016673 | chr17:57902814-57903248 | chrX:101914784-101914936 |
| chr3:170138195-170138413 | chr8:129053712-129054462 | chr11:94480491-94481212 | chr17:57906379-57907270 | chrX:106595212-106595632 |
| chr3:17913596-17913897 | chr8:129061512-129061955 | chr12:104658660-104659046 | chr17:60555037-60555507 | chrX:106595667-106595822 |
| chr3:182980917-182981737 | chr8:141598802-141599709 | chr12:104671230-104671489 | chr17:63639930-63640399 | chrX:110863259-110863845 |
| chr3:18580242-18580387 | chr8:142182442-142182863 | chr12:104693849-104694220 | chr17:64194072-64194322 | chrX:120337126-120337342 |
| chr3:194029385-194029774 | chr8:145002494-145002937 | chr12:104963305-104963849 | chr17:65220181-65220399 | chrX:125606277-125607115 |
| chr3:194040514-194040910 | chr8:145025737-145026802 | chr12:109120887-109121218 | chr17:66419326-66419636 | chrX:14262210-14262757 |
| chr3:25796078-25796406 | chr8:1693574-1694042 | chr12:109249958-109250678 | chr17:70499253-70499546 | chrX:25664451-25664736 |
| chr3:27674607-27674970 | chr8:1780723-1781665 | chr12:110867136-110867499 | chr17:70587266-70587776 | chrX:2845714-2845991 |
| chr3:39193999-39194528 | chr8:19266409-19266742 | chr12:119632995-119633391 | chr17:71064440-71064923 | chrX:39646201-39646652 |
| chr3:44040259-44040834 | chr8:19566953-19567512 | chr12:121268399-121268878 | chr17:75422359-75422648 | chrX:40693990-40694519 |
| chr3:44621262-44622718 | chr8:27730121-27730534 | chr12:125424077-125424667 | chr17:7737438-7738736 | chrX:40749971-40750359 |
| chr3:47704557-47705253 | chr8:29230784-29230978 | chr12:13025126-13025347 | chr17:7738822-7740614 | chrX:41182589-41182765 |
| chr3:5054958-5055470 | chr8:29522725-29523096 | chr12:132345340-132345736 | chr17:7745337-7745870 | chrX:43888590-43889088 |
| chr3:52080287-52080923 | chr8:29526859-29527039 | chr12:132970087-132970237 | chr17:7745915-7746944 | chrX:45702877-45703292 |
| chr3:53342432-53342824 | chr8:32603273-32603561 | chr12:132970628-132971029 | chr17:7746991-7747556 | chrX:46316672-46316948 |
| chr3:72977666-72978074 | chr8:35382808-35383133 | chr12:16553699-16554214 | chr17:79368790-79369424 | chrX:57421884-57422193 |
| chr3:73159773-73160145 | chr8:38895649-38895937 | chr12:16554253-16554721 | chr18:12288068-12288497 | chrX:67291116-67291490 |
| chr3:96336052-96336669 | chr8:43092918-43093442 | chr12:19260911-19261231 | chr18:18515990-18516559 | chrX:68004355-68004677 |
| chr3:98685741-98686133 | chr8:54884977-54885679 | chr12:20975224-20975562 | chr18:18517849-18518397 | chrX:70640504-70640913 |
| chr4:117519495-117520605 | chr8:57389900-57390701 | chr12:42825856-42826146 | chr18:29542474-29542964 | chrX:73507125-73507445 |
| chr4:122606332-122606884 | chr8:62620817-62621622 | chr12:46917975-46918280 | chr18:3624841-3625119 | chrX:85339971-85340972 |
| chr4:140970177-140970718 | chr8:62679636-62680132 | chr12:47033293-47033660 | chr18:5294371-5294839 | chrX:92479084-92479478 |
